# A Novel MicroRNA-Odorant Receptor Axis Governs Neural Progenitor Cell Proliferation in Zebrafish CNS

**DOI:** 10.64898/2026.05.26.727982

**Authors:** Samudra Gupta, Santosh Kumar Jana, Sukhendu Mandal, Arindam Biswas, Subhra Prakash Hui

## Abstract

Spinal cord injury causes irreversible neurological deficits in mammals; yet zebrafish achieve complete functional recovery through molecular mechanisms that remain poorly defined. In this study we emphasized on a critical miRNA-mediated regulation of Ependymo-radial glial (ERG) cell proliferation in zebrafish spinal cord. Using next-generation sequencing we constructed a spatiotemporal miRNA profile across multiple post-injury time points and identified dre-miR-N1 as a novel injury-responsive miRNA involved in ERG proliferation among several differentially expressed novel miRNAs. Fluorescent in situ hybridization confirmed its robust lesion-site expression and gain-of-function analysis demonstrated that dre-miR-N1 significantly impaired functional recovery. Target prediction and validation unexpectedly identified the odorant receptor gene *or42a1* as a high-confidence target and a combinatorial approach of miRNA gain-of-function and *or42a1* loss-of-function showed that dre-miR-N1 modulates the proliferative behaviour of *or42a1*-expressing ERG cells under both homeostatic and injury conditions. These findings uncover a previously unrecognized miRNA-odorant receptor axis governing injury-induced ERG cell expansion establishing a novel molecular framework for endogenous neural regeneration in zebrafish.

## 1. Introduction

Spinal cord injury (SCI) represents one of the most debilitating neurological conditions worldwide, disproportionately affecting young individuals and imposing profound physical, psychological, and socioeconomic burdens on patients and their families (Ahuja et al., 2017; Anjum et al., 2020; Guo, Redenski and Levenberg, 2021). Mechanistically, SCI is characterized by two temporally distinct phases: primary mechanical injury followed by a secondary injury cascade, the latter of which exacerbates tissue damage through local microenvironmental dysregulation and actively inhibits endogenous repair mechanisms (Lin et al., 2021; Wang et al., 2023). Considerable investigative effort has therefore shifted toward identifying the intrinsic molecular determinants governing post-injury microenvironmental remodeling and neural repair, with increasing recognition of the roles played by microtubule dynamics, epigenetic modulators, and non-coding RNAs (Chen et al., 2015; Li, Teng and Liu, 2016; Liu et al., 2020; Xu et al., 2021; Borger et al., 2022). Despite meaningful progress in these areas, effective repair of SCI remains a formidable global medical challenge, owing to the complexity of intertwined neuroprotective and regenerative mechanisms involved.

MicroRNAs (miRNAs) are small (∼22 nucleotide) non-coding RNAs that function as essential post-transcriptional regulators, primarily through base-pairing with complementary sequences in the 3’ untranslated region (3’UTR) of target mRNAs, leading to mRNA degradation or translational repression (Lu and Rothenberg, 2018; Galagali, 2023). As critical epigenetic modulators, miRNAs coordinate complex gene regulatory networks across diverse pathophysiological contexts. In mammalian SCI models, several miRNAs have been functionally implicated: mmu-miR-7a ameliorates injury by attenuating neuronal apoptosis and oxidative stress (Ding et al., 2020); mmu-miR-124-3p confers neuroprotection by suppressing neurotoxic microglial and astrocytic activation (Jiang et al., 2020); and the miR-219/miR-338 cluster promotes oligodendrocyte differentiation and myelination in vitro (Milbreta et al., 2019). These findings collectively underscore the capacity of individual miRNAs to orchestrate multiple facets of SCI pathophysiology.

In striking contrast to the limited regenerative capacity of mammals, adult zebrafish (*Danio rerio*) exhibit remarkable functional recovery within two weeks of complete spinal cord transection, positioning them as a uniquely valuable model for elucidating the cellular and molecular mechanisms underlying spinal cord regeneration (Becker and Becker, 2008; Hui, Dutta and Ghosh, 2010). Prior studies have implicated annotated miRNAs in zebrafish spinal cord regeneration through promotion of neural stem cell (NSC) proliferation and suppression of RhoA-mediated axonal regeneration inhibition (Yu et al., 2011; Li et al., 2021). However, the global miRNA landscape across discrete regenerative stages following SCI in zebrafish—and the functional contributions of novel, hitherto uncharacterized miRNAs to specific regenerative events—remain largely unexplored.

The present study addresses this knowledge gap by systematically characterizing the miRNA transcriptome across early regenerative time-points following complete spinal cord transection in adult zebrafish, identifying stage-specific differentially expressed novel miRNAs, and functionally dissecting the role of the top candidate, dre-miR-N1, in driving regenerative events. Strikingly, target prediction and functional validation identified the odorant receptor gene *or42a1*, canonically expressed in the nasal olfactory epithelium as a direct downstream effector of dre-miR-N1, exhibiting ectopic, injury-inducible expression in Sox2^+^ ependymo-radial glial (ERG) progenitor cells of the injured spinal cord. Loss- and gain-of-function experiments, validated by rescue assays, delineate a previously unrecognized dre-miR-N1/*or42a1* regulatory axis that governs ERG progenitor cell cycle entry and proliferative expansion during zebrafish spinal cord regeneration, revealing an unforeseen repurposing of a canonical sensory receptor gene in central nervous system repair.

## 2. Results

### 2.1. miRNA Biogenesis Is Indispensable for Functional Spinal Cord Regeneration

Pre-miRNAs are processed into mature miRNAs in the cytoplasm by the RNase III endonuclease Dicer. To determine whether global miRNA activity is required for zebrafish spinal cord regeneration, we locally inhibited Dicer using the pharmacological antagonist Cib3b delivered at the injury epicenter following complete spinal cord transection. Dose-dependent toxicity profiling in zebrafish embryos (up to 96 hpf) established a lethal concentration (LC50) and guided selection of 5 µM for local intra-spinal application, administered daily through 3 dpi (Fig.S1A–B). The efficacy of Dicer inhibition was confirmed by quantifying expression of the housekeeping miRNA dre-miR-92b-3p by stem-loop qRT-PCR; consistent with attenuated miRNA biogenesis, expression was uniformly low across uninjured and post-injury time points (6 hpi, 1, 3, and 7 dpi) without injury-induced modulation relative to DMSO-treated controls (Fig. S1C). Analysis of dre-miR-92b-3p levels in liver tissue of treated animals confirmed stable hepatic expression, demonstrating predominantly local Cib3b action at the injury site and minimal systemic exposure (Fig. S1D).

Following verification of Dicer inhibition and consequent global miRNA downregulation, we evaluated regenerative outcomes across three functionally relevant endpoints. GFAP immunostaining at 21 dpi revealed robust glial bridge formation at the lesion site in DMSO-treated controls, whereas Cib3b-treated animals exhibited a significant ∼31% reduction in glial bridging (p < 0.05; Fig. S2B-C). Axonal regrowth, assessed by anti-acetylated tubulin immunostaining at 30 dpi, was markedly impaired in Cib3b-treated zebrafish, which displayed disorganized axonal sprouts confined to the rostral and caudal stumps; approximately two-thirds of treated animals presented persistently severed stumps (Fig. S2D–E). Correspondingly, vehicle-treated controls fully restored pre-injury swimming capacity by 30 dpi, while Cib3b-treated animals exhibited significantly impeded functional recovery (p < 0.05; Fig. S2F–G). Collectively, these data establish that miRNA biogenesis is indispensable for glial bridging, axonal regrowth, and functional locomotor recovery following SCI in adult zebrafish.

### 2.2. Small RNA Sequencing Identifies Novel miRNAs Across Early Regenerative Stages

To comprehensively profile the miRNA transcriptome during post-injury regeneration, five small RNA sequencing libraries from adult zebrafish spinal cord (n = 6 per time point) were pooled to maximize read depth for novel miRNA discovery. The combined dataset comprised 250,124,276 raw reads (Fig. 1A), satisfying the stringent read-depth requirements established for reliable novel miRNA identification (Dhahbi et al., 2011; Szakats, McAtamney and Wilson, 2024). miRDeep2 analysis identified 2,586 miRNAs expressed in the regenerating spinal cord, validating the biological representativeness of the dataset. Applying the established miRDeep2 score threshold (≥2 × 10^2^)(McCreight et al., 2017), the novel discovery module predicted 189 candidate miRNAs (Fig. 1A).

**Figure 1:**
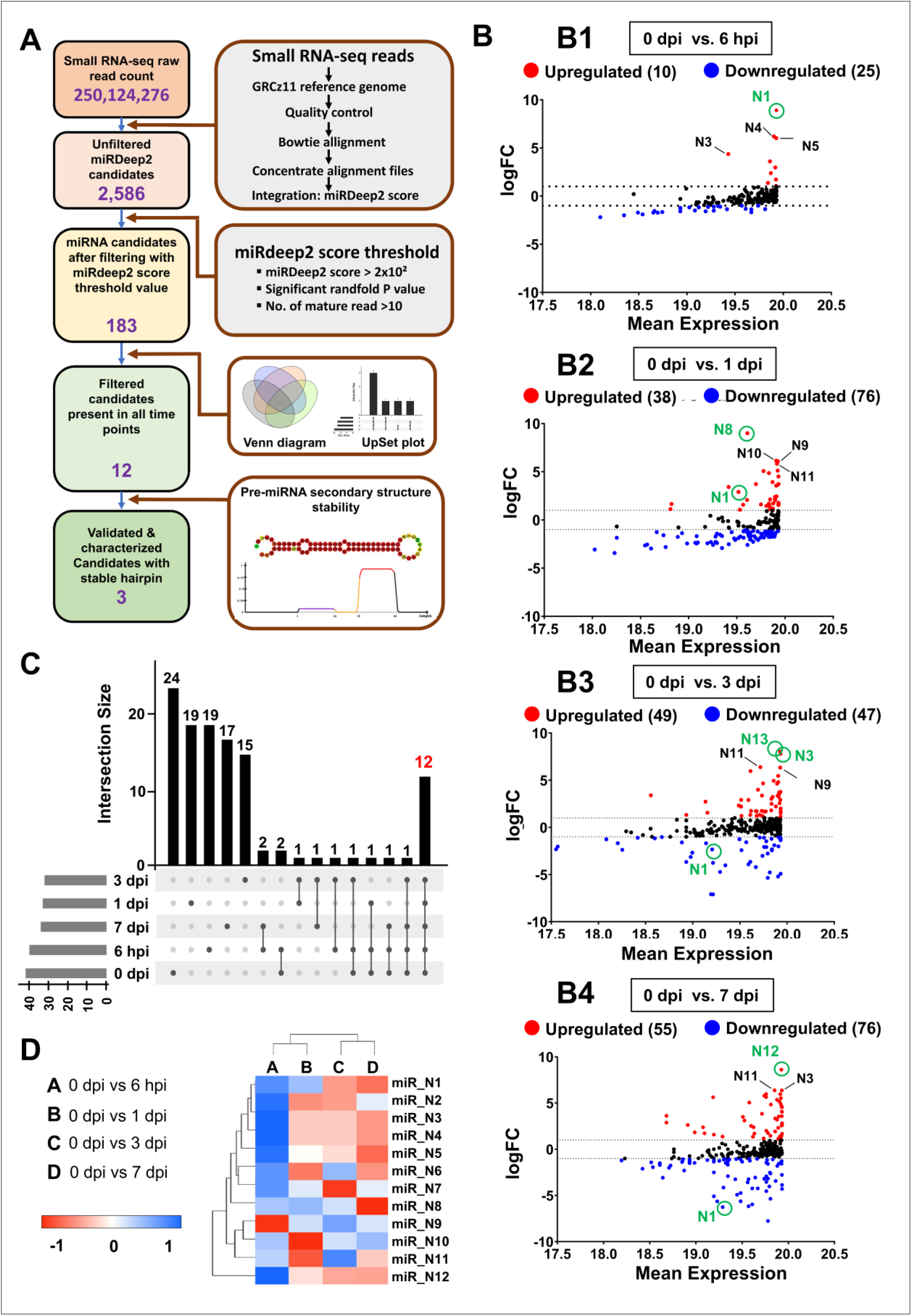
Novel miRNA discovery pipeline and differential expression of novel miRNAs during regeneration. (A) Flowchart showing the Analyses and number of novel miRNAs at each step in the discovery pipeline. (B) MA plots showing differential expression of miRdeep2 filtered candidates among comparisons of 0dpi vs 6 hpi (B1), 0dpi vs 1 dpi (B2), 0 dpi vs 3 dpi (B3), and 0 dpi vs 7 dpi (B4). Green circle marks the significant novel miRNA in each time course. (C) UpSet plot diagram of differentially expressed miRNAs (log2 (fold-change) > 0.5, p < 0.05) showing the total set size and overlaps between the 183 novel miRNAs and those found in all time points. The y-axis indicates the number of commonly occurring novel miRNAs identified in each time point. The intersecting miRNA datasets are presented as shaded circles connected by solid lines in the lower panel. Common miRNAs intersected by all time points marked in red.(D) Heatmap expression of 12 differentially expressed novel miRNAs among comparisons of 0dpi vs 6 hpi, 0dpi vs 1 dpi, 0 dpi vs 3 dpi, and 0 dpi vs 7 dpi. All these 12 novel miRNAs are differentially expressed in all groups [log2 (fold-change) >1, p < 0.01].

### 2.3. Identification and Characterization of Thermodynamically Stable Three Novel miRNAs During the Early Regenerative Response

To minimize false positives, the 189 candidate novel miRNAs were cross-compared across all five experimental time points (0 dpi, 6 hpi, 1 dpi, 3 dpi, and 7 dpi), retaining only those consistently detected under all conditions. This stringent intersection yielded 12 novel miRNA candidates common to all time points (Fig. 1A, Supplementary Table S1). These candidates were further subjected to thermodynamic validation using RNA secondary structure prediction; stable hairpin conformations with low minimum free energy (MFE) values were required, reflecting greater structural stability of the predicted pre-miRNA hairpin (Lorenz, Bernhart and Siederdissen, 2011). Applying criteria established earlier (Zayed, Qi and Peng, 2019), RNAfold analysis of the 12 filtered candidates identified three thermodynamically optimal stem-loop structures, designated dre-miR-N1, dre-miR-N2, and dre-miR-N3.

### 2.4. Differential Expression Analysis Reveals Extensive Time-Dependent miRNA Regulation During Regeneration

Differential expression analysis relative to uninjured spinal cord revealed extensive, time-dependent miRNA regulation: 183 novel miRNAs were differentially expressed at 6 hpi (10 upregulated, 25 downregulated), 176 at 1 dpi (38 upregulated, 76 downregulated), 326 at 3 dpi (49 upregulated, 47 downregulated), and 321 at 7 dpi (55 upregulated, 76 downregulated) (Fig. 1B). UpSet plot analysis of the 189 candidates confirmed 12 novel miRNAs with consistent presence across all experimental time points and the highest coefficients of variation (Fig. 1C). Hierarchical cluster analysis of these 12 differentially expressed novel miRNAs revealed distinct temporal expression clusters (Fig. 1D), from which the three thermodynamically stable candidates were advanced for qRT-PCR validation.

dre-miR-N1 exhibited the highest miRDeep2 score (4,186.4), a significant randfold p-value, and a mature read count of 4,394 reads, with no homology to any previously annotated miRNA. qRT-PCR confirmed high basal expression in uninjured spinal cord that was maintained at 6 hpi, followed by a progressive and statistically significant decline from 1 dpi through 7 dpi (p < 0.01; Fig. 2A). dre-miR-N2 displayed a high miRDeep2 score (2,241), a significant randfold p-value, and a mature read count of 953, with abundant basal expression declining progressively through 3 dpi to near-undetectable levels, before rebounding substantially at 7 dpi to exceed uninjured baseline levels (Fig. 2A). dre-miR-N3 exhibited consistently high expression across all time points including uninjured condition with a very low level of expression at 7 dpi (Fig. 2A).

**Figure 2:**
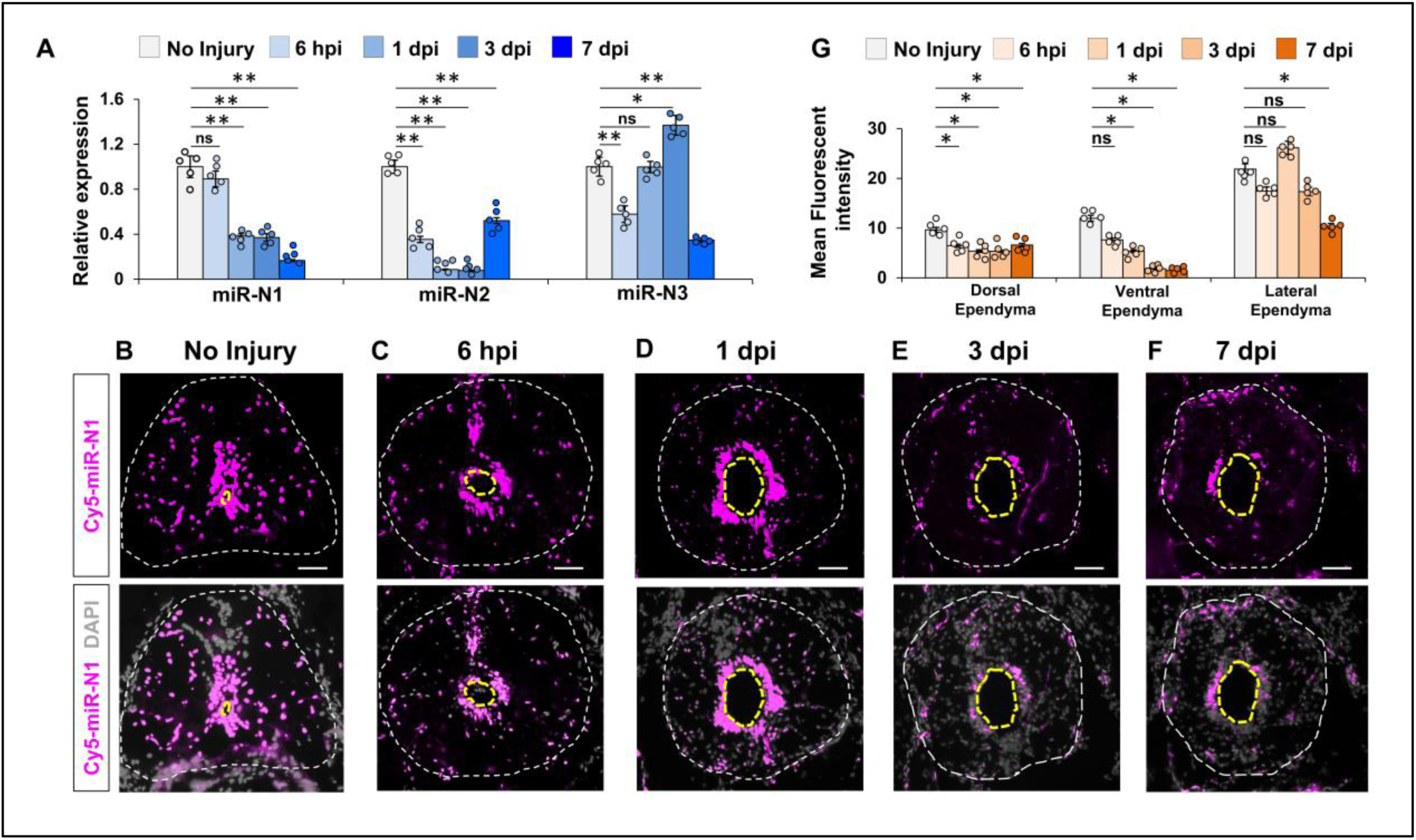
Spatio-temporal expression profile of dre-miR-N1 in regenerating spinal cord. (A) qRT-PCR analysis of three most stable novel miRNAs time-course expression in regenerating spinal cord. Data represented as Mean±SEM, n=5; **p < 0.01; Mann-Whitney U test. (B-F) Spatio-temporal expression of novel dre-miR-N1 using Cy5-labelled LNA-probe mediated Fluorescent in situ hybridization (FISH) followed by DAPI staining in regenerating spinal cord (transverse section). The white-dotted line marked border of the spinal cord and yellow-dotted line marked the ependymal canal; Magnification, 10x; Scale bar, 50µm. (G) Quantification of mean fluorescent intensity of Cy5-labelled LNA probe signal in regions dorsal, ventral, and lateral to the ependymal canal. Data represented as Mean±SEM, n=5; **p < 0.01; Mann-Whitney U test.

### 2.5. dre-miR-N1 Is Selected as the Priority Candidate for Functional Investigation

Among the three validated candidates, dre-miR-N1 was designated the primary target for functional investigation based on its superior bioinformatic characteristics: the highest miRDeep2 score, the greatest mature read count (4,394 reads), and the most thermodynamically favorable pre-miRNA structure (MFE: −44.50 kcal/mol). Its expression profile—high during the uninjured and acute post-injury phases, with progressive downregulation through the proliferative regenerative phase is consistent with a miRNA functioning as a regulatory brake on ERG progenitor activation that is subsequently attenuated to permit downstream regenerative events. The pronounced and consistent expression dynamics of dre-miR-N1, substantiated by both small RNA sequencing and qRT-PCR, provided the strongest rationale for prioritizing this candidate in mechanistic dissection of post-injury miRNA function.

### 2.6. Spatio-Temporal Expression of dre-miR-N1 Is Enriched in Sox2^+^ ERG Progenitor Cells

To delineate the injury-induced spatiotemporal expression dynamics of dre-miR-N1, fluorescent in situ hybridization (FISH) was performed on cryosections from each regenerative time point using a Cy5-labeled locked nucleic acid (LNA) probe (Fig. 2B–F). In uninjured spinal cord, dre-miR-N1 signal was broadly distributed across white and gray matter (Fig. 2B). At 6 hpi, signal remained abundant in both compartments with notable attenuation in the parenchyma (Fig. 2C). By 1 dpi, expression became markedly enriched around the ependymal canal while continuing to diminish in the parenchyma (Fig. 2D). At 3 and 7 dpi, global expression declined further, but remained concentrated at the ependymal canal (Fig. 2E–F).

Quantitative fluorescence intensity analysis across three ependymal zones—dorsal, lateral, and ventral—revealed region-specific dynamics (Fig. 2G). A significant and progressive reduction in dre-miR-N1 expression was observed in the ventral ependyma beginning at 6 hpi, with levels becoming nearly undetectable by 7 dpi (p < 0.05; Fig. 2G). In contrast, lateral ependymal expression was consistently elevated throughout the regenerative time course, peaking at 1 dpi and thereby distinguishing this compartment from both dorsal and ventral regions (Fig. 2G). Co-immunostaining of FISH-labelled sections with anti-GFAP and anti-Sox2 antibodies revealed colocalization of dre-miR-N1 signal with Sox2^+^ and GFAP^+^ ependymal cells in both uninjured and 7 dpi conditions, with no parenchymal colocalization (Fig. S4A–B). These findings indicate that dre-miR-N1 is predominantly expressed by Sox2^+^ ERG progenitor cells, implicating it as a regulator of their activity during regeneration.

### 2.7. Gain-of-Function dre-miR-N1 Overexpression Impairs Endpoints of Spinal Cord Regeneration

A synthetic dre-miR-N1 mimic was produced by in vitro transcription (Sata et al., 2025). Dose-optimization via intra-spinal delivery in a gel-foam vehicle established that the 2 µg dose produced a 4.75-fold increase in total miRNA without perturbing expression of housekeeping miRNAs (dre-miR-92b-3p, dre-miR-206-3p) or the reference mRNA *elf1a* (*eef1a1l1)*, while significantly downregulating *or42a1* mRNA (p < 0.01; Fig. S3B–D), confirming dose-specific and physiologically plausible mimic activity. Intra-spinal dre-miR-N1 mimic delivery was performed daily from transection through 7 dpi (Fig. 3A).

**Figure 3:**
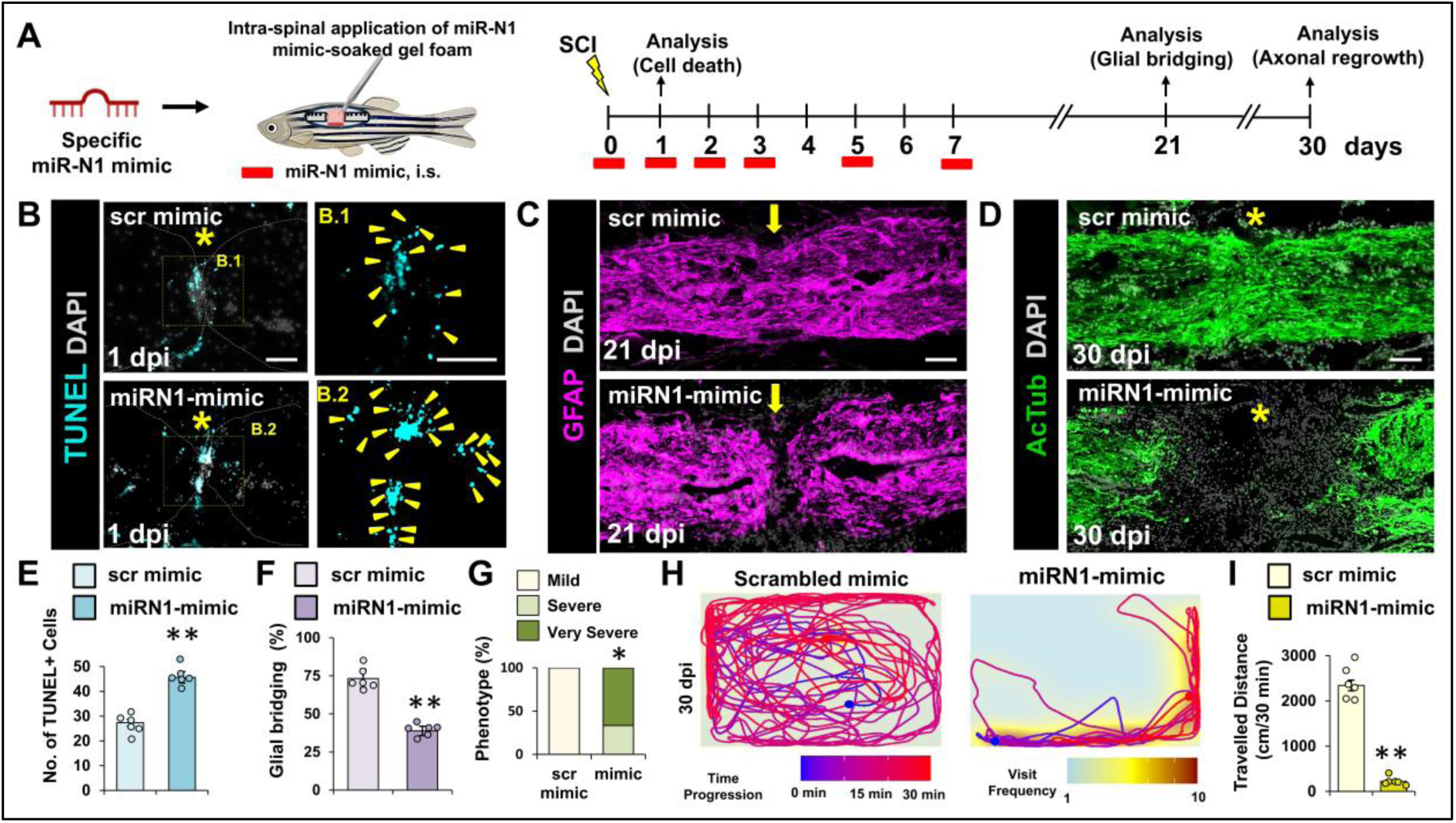
Overexpression of novel dre-miR-N1 impedes the phenotype recovery after spinal cord injury. (A) Experimental intervention involving novel dre-miR-N1 mimic treatment. i.s. intra-spinal injection. (B) Immunohistochemical staining of spinal cord sections after 1 dpi from scrambled mimic-treated and dre-miR-N1 mimic-treated fish showing TUNEL-positive cells at the injury epicentre. Insets show magnification of the demarcated region; Scale bar, 50µm. (C) Immunohistochemical staining of spinal cord sections after 21 dpi from scrambled mimic-treated and dre-miR-N1 mimic-treated fish showing glial bridge formation. (D)Immunohistochemical staining of spinal cord sections after 30 dpi from scrambled mimic-treated and dre-miR-N1 mimic-treated fish showing axonal projections. (E) Quantification of TUNEL-positive cells at the injury epicentre in (B) (N=6). (F)Quantification of glial bridging in (C) (N=6). (G) Quantification of regeneration in (D) (N=5). (H)Swim tracking of individual animals after 30 dpi from scrambled mimic-treated and dre-miR-N1 mimic-treated experimental groups (N=8). (I)Quantification of total distance moved in (H). Yellow arrow-heads marked the TUNEL-positive cells; Yellow arrow indicates glial bridging; Asterisks indicate injury epicenter. Confocal projections of z stacks are shown (C, D). *p < 0.05; **p < 0.01; Fisher’s exact test (G) and Mann-Whitney U test (E, F, I). Magnification, 10x (B, C, D), 20X (B1, B2); Scale bars, 50 µm (B, C, D) and 0.1 m (H).

TUNEL assay at 1 dpi revealed a significant increase in apoptotic cells at the injury epicenter in mimic-treated animals relative to scrambled mimic controls (p < 0.01; Fig. 3B, 3E), implicating dre-miR-N1 in early post-injury cell survival regulation. At 21 dpi, GFAP immunostaining demonstrated a ∼44% reduction in glial bridge area in mimic-treated animals (p < 0.01; Fig. 3C, 3F). Axonal regrowth assessed by anti-acetylated tubulin immunostaining at 30 dpi revealed that the majority of mimic-treated animals displayed disorganized axonal sprouts with persistent rostro-caudal stump discontinuity (Fig. 3D; Severe and Very Severe categories in Fig. 3G), contrasting with near-complete regeneration in scrambled mimic controls. Locomotor analysis at 30 dpi demonstrated a ∼68% reduction in swimming ability in dre-miR-N1 mimic-treated animals (p < 0.05), quantified across total distance travelled, mean velocity, mean swim speed, and burst frequency (Fig. 3H–I; Fig. S4A–C). These findings collectively establish that sustained dre-miR-N1 overexpression impairs cell survival, glial bridge formation, axonal regrowth, and functional locomotor recovery

### 2.8. *In Silico* Target Prediction and Biochemical Validation Identify *or42a1* as the Primary Effector of dre-miR-N1

Seed sequence homology analysis identified dre-miR-153c as the closest annotated miRNA match to dre-miR-N1 (E-value = 0.001), with additional matches to four dre-miR-153 family members and one murine and one human homolog at progressively higher E-values (Fig. 4A). Integrated target prediction using TargetScan (context score < −0.2; 279 predicted genes) and DIANA microT-CDS v.5.0 (miTG score > 0.7; 289 predicted genes) identified 236 high-confidence overlapping targets (Fig. 4B). Gene Ontology (GO) analysis of these 236 candidates yielded 26 significantly enriched terms (p < 0.05), with 8 upregulated genes linked to overrepresented terms associated with early regenerative processes (Fig. 4C). Three genes—*nodal-related 1* (*ndr1*), *teashirt zinc finger homeobox 1* (*tshz1*), and *odorant receptor 42a1* (*or42a1*)—emerged as top candidates based on GO enrichment breadth. *or42a1* was associated with the largest number of significant GO terms (n = 13), spanning cell migration (GO:0030334, GO:0031346), cellular differentiation (GO:0045597, GO:0032535, GO:0048857), and granulocyte chemotaxis (GO:0090022), while retaining association with its canonical olfactory function (GO:0050911).

**Figure 4:**
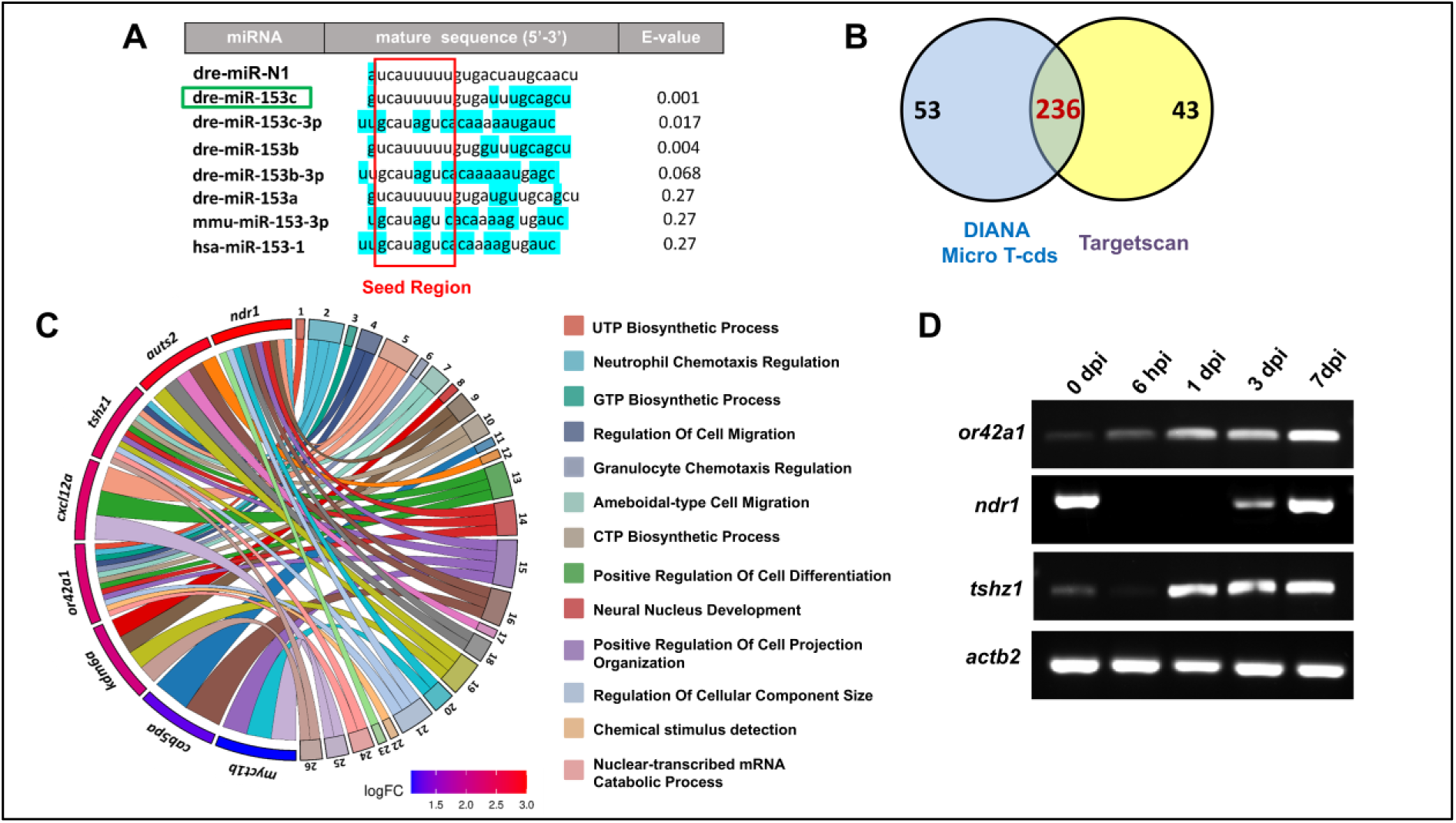
Functional target prediction for novel dre-miR-N1. (A) The miRNA seed sequence family analysis identified known annotated dre-miR-153c in zebrafish and other species with similar seed sequences using Blastn. They were ranked with the lowest E-values. Red box marked the seed sequence in the mature sequence of all the miRNAs. Blue highlighted bases marked the base-pair mismatch. Green box indicates the annotated miRNA with lowest E-value and no base-pair mismatch. (B) Target gene identification from two tools (DIANA Micro T-cds and Targetscan) for the annotated miRNA, dre-miR-153c, with lowest E-value. The Venn diagram indicates the common genes identified by both tools. (C) GO Chord showing the significant (p-adj<0.05) and top upregulated (log2FC>1) genes after gProfiler GO enrichment analysis for the 236 predicted target genes. GO terms associated with *or42a1* mentioned beside the GO chord. (D) Gel images of RT-PCR expression of *or42a1*, *tshz1*, and *ndr1* during regeneration time course in zebrafish spinal cord.

Bioinformatic duplex modeling using RNAhybrid (Rehmsmeier et al., 2004) predicted a thermodynamically stable complex between the 8-mer seed region of dre-miR-N1 and the *or42a1* 3’UTR, featuring perfect Watson-Crick complementarity and an MFE of −22.2 kcal/mol (Fig. 5A), compared to the less stable duplexes predicted for *tshz1* (MFE: −15.97 kcal/mol; one mismatch; Fig. 5B) and *ndr1* (MFE: −8.72 kcal/mol; one mismatch; Fig. 5C). RNA electrophoretic mobility shift assay (EMSA) confirmed concentration-dependent complex formation between dre-miR-N1 and *or42a1* 3’UTR RNA at 10 and 20 µM (Fig. 5D), with superior binding affinity relative to *tshz1* and *ndr1*, as reflected by progressive reduction of unbound miRNA fractions (Fig. 5E–G). Luciferase reporter assays in HEK293T cells demonstrated significant repression of wildtype *or42a1* 3’UTR-driven luciferase activity by dre-miR-N1 mimic (p < 0.01), with no significant effect on the mutant 3’UTR construct (Fig. 5H), unambiguously confirming *or42a1* as a direct and functional transcriptional target of dre-miR-N1

**Figure 5:**
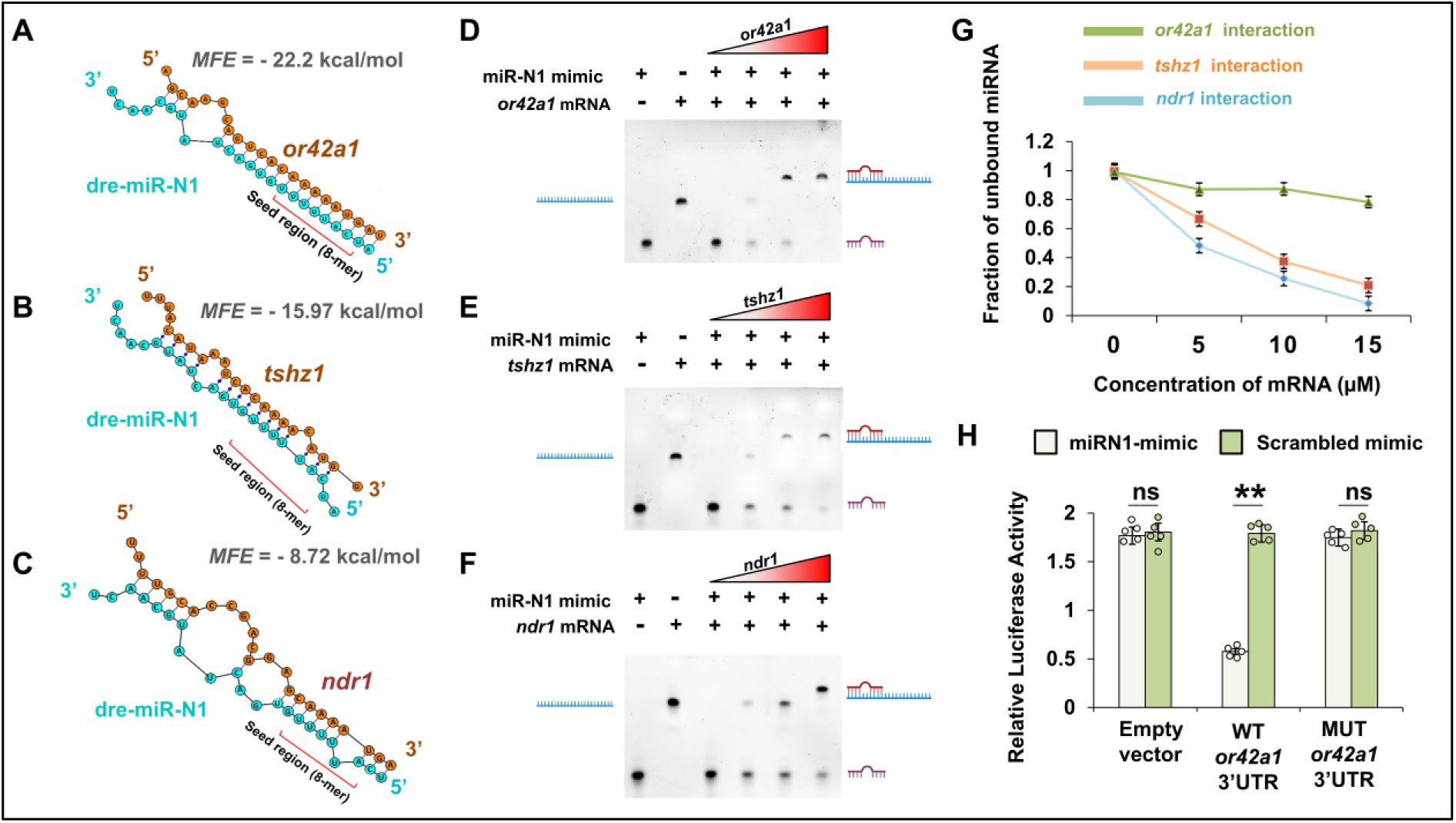
in silico target prediction and functional validation for novel dre-miR-N1. (A-C) RNAhybrid results showing the binding between 3’ UTR region of three target mRNAs (*or42a1, tshz1, and ndr1*) and the seed region of the candidate dre-miR-N1 and the minimum free energy (MFE) for the hybridization. (D-F) Validation of miRNA-mRNA binding by RNA-EMSA method between the three target genes and the novel dre-miR-N1. dre-miR-N1 mimic and target gene mRNA were synthesized by in vitro transcription and used for validation of miRNA-mRNA binding. Lane 1 corresponds to the candidate dre-miR-N1 alone, Lane 2 corresponds to the 3’UTR of the target mRNA alone; Lane 3,4,5 and 6 correspond to the incubation of the dre-miR-N1 mimic with increasing concentrations (0 µM,5 µM, 10µM, and 20 µM) of 3’UTR of target mRNA of *or42a1* (D), tshz1 (E), and, ndr1 (F), respectively. Schematic indication of miRNA-mRNA complex, 3’UTR of target mRNA, and free unbound miRNA was marked beside the gel images. (G) Plot showing the fraction of unbound miRNA at varying mRNA concentrations. The fraction of unbound miRNA in case of *or42a1*, *tshz1* and *ndr1* were calculated from the densiometric analysis (N=3) of the free unbound miRNA bands in D, E and F. For control in densiometric calculation, only miRNA was run in four lanes in the same gel parallelly. (H) Relative Luciferase activity (RLU) of sicheck2 vector and sicheck2 plasmid with wildtype *or42a1* 3’UTR and mutated *or42a1* 3’UTR inserts co-transfected with scrambled mimic and miRN1 mimic in HEK293T cells. Data represented as Mean ± SEM (N=3). ** P<0.01; ns, non-significant; Mann-Whitney U test.

### 2.9. *or42a1* Exhibits Ectopic Injury-Inducible Expression and Promotes ERG Progenitor Proliferation

*or42a1* encodes a G protein-coupled receptor canonically expressed in the nasal olfactory epithelium for odorant detection (Barth, Dugas and Ngai, 1997; Franco et al., 2025). qRT-PCR analysis revealed low *or42a1* transcript levels in uninjured and 6 hpi spinal cord, followed by significant upregulation at 1 dpi, peaking at 3 and 7 dpi (p < 0.01; Fig. 6A)—a temporal expression profile reciprocal to that of dre-miR-N1. In situ hybridization (ISH) confirmed injury-induced *or42a1* transcript accumulation adjacent to the ependymal canal from 1 dpi onward, with robust expression in ependymal and pericanal cells at 3 dpi (Fig. 6B–D). Co-immunostaining with anti-Sox2 antibody identified *or42a1*^+^ cells as Sox2^+^ ERG progenitor cells (Fig. 6E). In contrast, *tshz1* and *ndr1* expression patterns were not consistent with dre-miR-N1-mediated regulation (Fig. 4D), further prioritizing *or42a1* as the primary functional effector.

**Figure 6:**
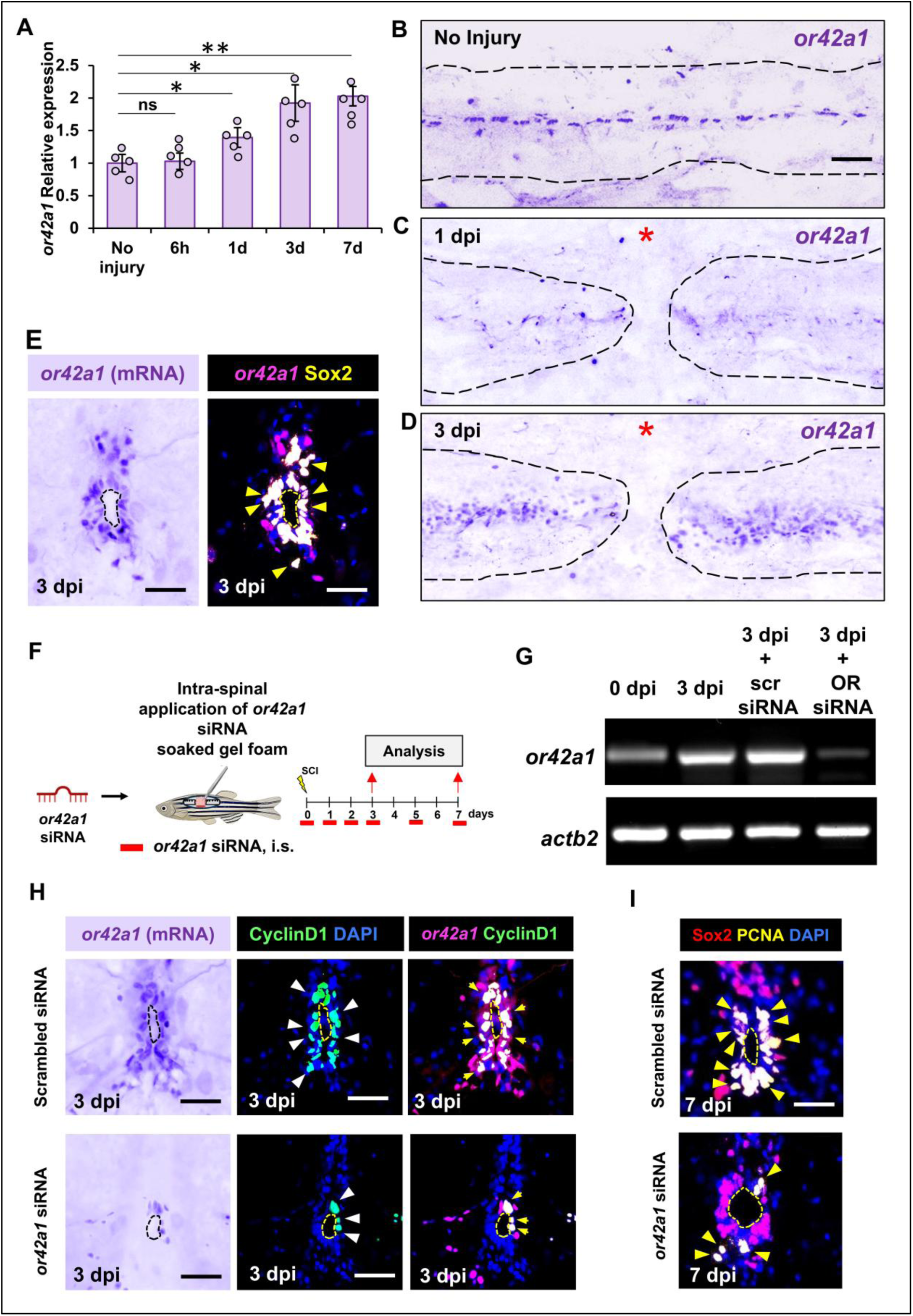
*or42a1* regulates proliferation of ERG progenitor cells during regeneration. (A) qRT-PCR analysis of *or42a1* time-course expression in regenerating spinal cord. Data represented as Mean±SEM, n=5; *p<0.05, **p < 0.01, ns, non-significant; Mann-Whitney U test. (B-D) In situ hybridization for *or42a1* in uninjured (B),1 dpi (C), and 3 dpi (D) spinal cord longitudinal sections. Dotted lines delineate the spinal cord. Asterisks mark the injury epicenter. (E) In situ hybridization for *or42a1* in 3 dpi spinal cord in combination with immunofluorescent staining for Sox2. Digitally merged image from in situ and immunostaining signals are shown on the right with Sox2^+^ cells in yellow, nuclei in blue and in situ signals digitally converted into magenta. Yellow arrowheads mark the Sox2^+^ cells coexpressing *or42a1* signals. Yellow dotted line marked the ependymal canal. (F) Experimental intervention involving *or42a1* siRNA treatment. i.s. intra-spinal injection. (G) Representative gel images showing the expression of *or42a1* transcript at 3 dpi including uninjured condition and in both scrambled siRNA and *or42a1* siRNA treated condition. (H) In situ hybridization for *or42a1* in 3 dpi injured spinal cord in combination with immunostaining for CyclinD1 marker in both scrambled siRNA and *or42a1* siRNA treated condition. Digitally merged images from in situ and immunostaining signals are shown on the right with CyclinD1^+^ cells in green, nuclei in blue and in situ signals digitally converted into magenta. White arrowheads marked the CyclinD1^+^ cells and yellow arrows marked the CyclinD1^+^ cells coexpressing *or42a1* transcript. Yellow and black dotted line delineate the ependymal canal. Magnification, 20x; Scale bar, 50µm. (I) Immunohistochemical staining of 7 dpi injured spinal cord in scrambled siRNA-treated and *or42a1* siRNA-treated experimental groups showing expression of Sox2 and PCNA. Yellow arrowheads marked the Sox2^+^PCNA^+^ proliferating NPCs. Magnification, 20x; Scale bar, 50µm.

To assess the functional role of *or42a1*, intra-spinal siRNA-mediated knockdown was performed following transection (Fig. 6F), with qRT-PCR and ISH confirming significant *or42a1* mRNA reduction at 3 dpi relative to scrambled siRNA controls (Fig. 6G–H). CyclinD1 immunostaining—marking G1-phase cells primed for proliferative entry—revealed a marked decrease in *or42a1*^+^/CyclinD1^+^ double-positive cells in siRNA-treated cords at 3 dpi (Fig. 6H), indicating impaired G1/S transition. Quantification of Sox2^+^/PCNA^+^ proliferating ERG progenitors at 7 dpi demonstrated a significant reduction following *or42a1* knockdown compared to scrambled controls (Fig. 6I; Fig. S5D), establishing *or42a1* as a positive regulator of ERG progenitor proliferation.

To confirm dre-miR-N1-mediated *or42a1* regulation in vivo, gain-of-function mimic delivery was performed and *or42a1* transcript levels assessed at 3 dpi. qRT-PCR and ISH both demonstrated significant *or42a1* mRNA suppression in mimic-treated animals (Fig. 7B–C). CyclinD1^+^/*or42a1*^+^ double-positive cells at the ependymal canal were markedly reduced (p < 0.01; Fig. 7C), and Sox2^+^/PCNA^+^ proliferating ERG progenitors were significantly diminished at 7 dpi (p < 0.01; Fig. 7D; Fig. S5E). Critically, rescue experiments involving co-delivery of dre-miR-N1 mimic with an *or42a1* coding-sequence (CDS) mimic—lacking the miRNA binding site and thus resistant to dre-miR-N1-mediated repression—restored Sox2^+^/PCNA^+^ ERG progenitor numbers to approximately twofold above the dre-miR-N1 mimic + scrambled mimic control at 7 dpi (p < 0.01; Fig. 7E; Fig. S5F). These findings conclusively establish that dre-miR-N1 suppresses ERG progenitor proliferation via direct repression of *or42a1*, defining the dre-miR-N1/*or42a1* regulatory axis as a functionally essential mechanism for progenitor-driven spinal cord regeneration in adult zebrafish.

**Figure 7:**
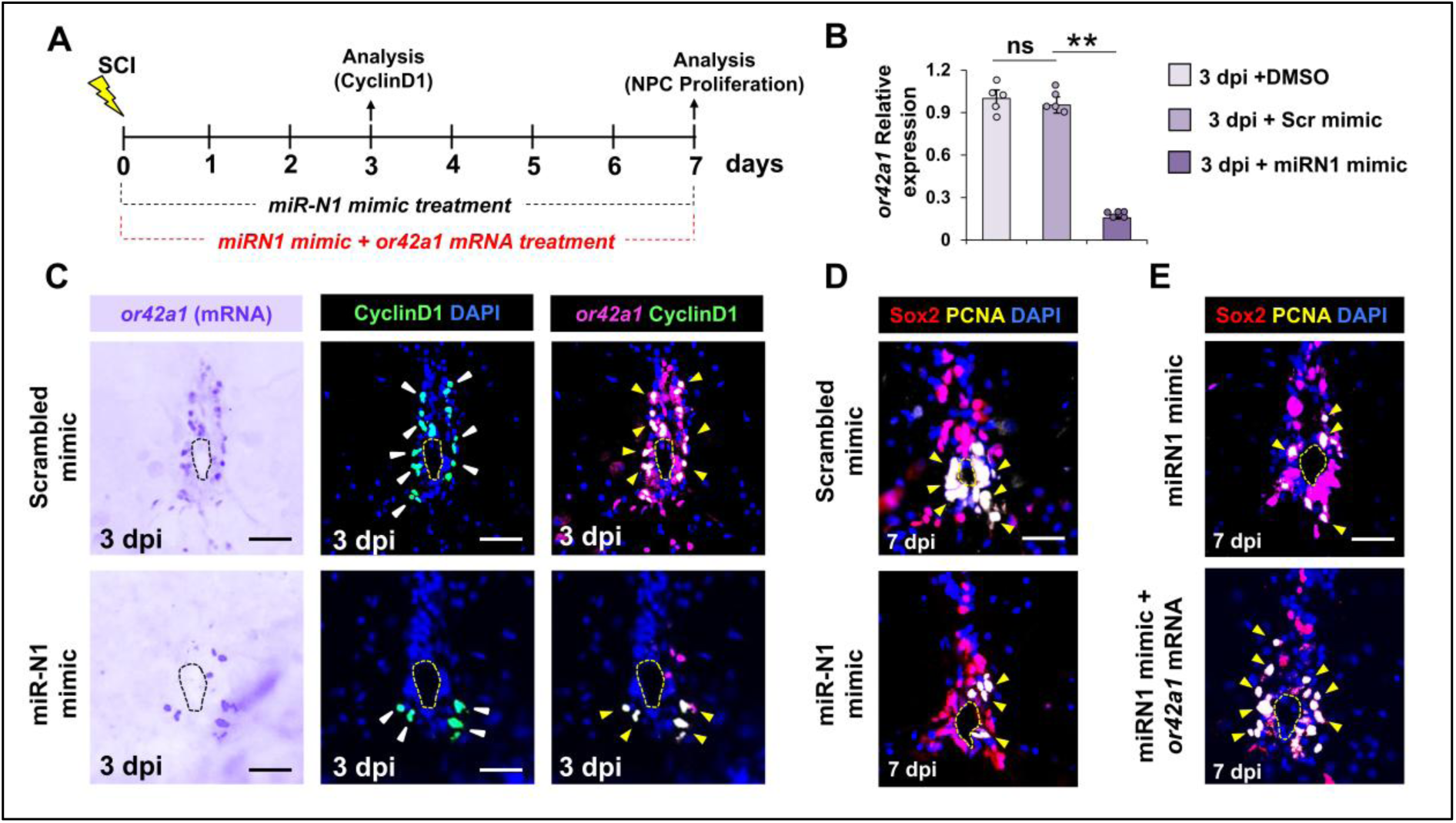
Novel dre-miR-N1 modulates ERG progenitor proliferation by regulating *or42a1* expression during regeneration. (A) Experimental intervention involving dre-miR-N1 mimic treatment. i.s. intra-spinal injection. (B) qRT-PCR analysis of *or42a1* at 3 dpi in DMSO-treated, scrambled-mimic treated and miRN1 specific mimic treated group. Data represented as Mean±SEM, n=5; *p<0.05, **p < 0.01, ns, non-significant; Mann-Whitney U test. (C) In situ hybridization for *or42a1* in 3 dpi injured spinal cord in combination with immunostaining for CyclinD1 in both scrambled mimic-treated control and dre-miR-N1 mimic treated condition. Digitally merged images from in situ and immunostaining signals are shown on the right with CyclinD1^+^ cells in green, nuclei in blue and in situ signals digitally converted into magenta. White arrow heads marked the CyclinD1^+^ cells and yellow arrows marked the CyclinD1^+^ cells coexpressing *or42a1* transcript. (D) Immunohistochemical staining of 7 dpi injured spinal cord in both scrambled mimic-treated control and dre-miR-N1 mimic treated experimental groups showing expression of Sox2 and PCNA. Yellow arrowheads marked the Sox2^+^PCNA^+^ proliferating NPCs. (E) Immunohistochemical staining of 7 dpi injured spinal cord in both dre-miR-N1 mimic treated and or42a1 CDS mRNA treated rescue experimental groups showing rescued expression of Sox2 and PCNA. Yellow arrowheads marked the Sox2^+^PCNA^+^ proliferating NPCs. Magnification, 20x; Scale bar, 50µm.

## 3. Discussion

The capacity for robust spinal cord regeneration in zebrafish offers a compelling framework for identifying conserved molecular mechanisms that may inform strategies to promote repair in non-regenerative species. The present study demonstrates that miRNA-mediated post-transcriptional regulation constitutes a pivotal epigenetic layer governing early regenerative events following spinal cord transection in adult zebrafish. By integrating pharmacological Dicer inhibition, high-throughput small RNA sequencing, bioinformatic target identification, and multidimensional gain- and loss-of-function experiments, we identify a novel miRNA, dre-miR-N1 and delineate its functional interaction with the ectopically expressed odorant receptor *or42a1* as a critical regulatory axis governing ERG progenitor proliferation during spinal cord regeneration.

The role of miRNAs in SCI pathophysiology spanning neuroprotection, neuroinflammation resolution, and axonal regeneration has been established primarily in mammalian model systems (Milbreta et al., 2019; Ding et al., 2020; Jiang et al., 2020). The contribution of epigenetic mechanisms to zebrafish spinal cord regeneration was recently substantiated by our own prior work demonstrating that Sirtuin1, a class III histone deacetylase, regulates axonal regeneration via modulation of the Hippo signaling pathway (Gupta and Hui, 2025). The present study extends this epigenetic regulatory framework to miRNA-mediated control, complementing prior investigations of annotated miRNAs in zebrafish spinal cord regeneration (Yu et al., 2011; Huang et al., 2017; Shen et al., 2024), while uniquely demonstrating the functional significance of novel, previously uncharacterized miRNAs in orchestrating discrete regenerative events.

Pharmacological inhibition of Dicer-mediated miRNA biogenesis using Cib3b produced a comprehensive disruption of regenerative outcomes impaired glial bridging, deficient axonal regrowth, and attenuated locomotor recovery establishing a global requirement for miRNA activity in functional spinal cord repair. The stability of dre-miR-92b-3p in liver tissue of treated animals confirmed predominantly local Cib3b action, minimizing the confound of systemic miRNA suppression. These observations provided the biological rationale for systematic transcriptomic characterization of the post-injury miRNA landscape through small RNA sequencing.

Small RNA sequencing of 250 million reads across five temporally defined stages spanning the transition from uninjured cord to peak NPC proliferation yielded a comprehensive miRNA transcriptome of the regenerating zebrafish spinal cord. The application of stringent miRDeep2 score thresholds, cross-time point consistency filtering, and thermodynamic validation via MFE-based secondary structure prediction yielded three robust novel miRNA candidates. Among these, dre-miR-N1 emerged as the functionally most compelling candidate, distinguished by its superior miRDeep2 score, highest mature read count, and most stable pre-miRNA secondary structure. Its expression profile high basally and in the acute injury phase, with progressive downregulation through the proliferative phase is characteristic of a miRNA functioning as a molecular brake on regenerative activation, analogous to paradigms established for miRNAs that maintain progenitor quiescence in diverse regenerative contexts.

The spatio-temporal localization of dre-miR-N1 to Sox2^+^ ERG progenitor cells surrounding the ependymal canal carries important mechanistic implications. Prior studies have established that Ctgfa^+^/GFAP^+^ ventral ependymal cells undergo epithelial-to-mesenchymal transition (EMT) prior to entering the proliferative cycle to form the glial bridge (Mokalled et al., 2016; Klatt Shaw et al., 2021), and that Olig2-expressing ERG cells in the ventrolateral domain initiate neuronal differentiation programs during regeneration (Reimer et al., 2009; Kuscha et al., 2012). Our observation that dre-miR-N1 expression is selectively diminished in the ventral ependyma with regenerative progression while being sustained in lateral ependymal and parenchymal compartments indicates that its downregulation in ventral Sox2^+^ ERGs is a prerequisite for EMT initiation and subsequent proliferative expansion. The sustained expression in lateral and parenchymal compartments is consistent with minimal involvement of dre-miR-N1 in neuronal lineage specification, situating its primary function in the regulation of progenitor activation rather than terminal differentiation.

The identification of *or42a1* as the primary functional target of dre-miR-N1 constitutes a significant conceptual advance. Odorant receptors (ORs) are G protein-coupled receptors conventionally ascribed exclusive roles in olfactory detection of volatile compounds in the nasal epithelium (Patel and Peralta-Yahya, 2023). However, a growing body of evidence documents ectopic OR expression and non-canonical functions in diverse non-olfactory tissues including heart, skin, sperm, intestine, and multiple brain regions (Sato, Miyasaka and Yoshihara, 2007; Oh, 2018). In the zebrafish genome, approximately 140 OR genes are expressed from 24 hours post-fertilization onward (Barth, Dugas and Ngai, 1997; Argo, Weth and Korsching, 2003; Sato, Miyasaka and Yoshihara, 2007), and recent studies have demonstrated OR expression in extra-olfactory tissues including muscle, eye, pharynx, and several brain regions such as the optic tectum, hypothalamus, habenula, and brainstem (Jundi et al., 2023). Our study reports for the first time the ectopic, injury-inducible expression of *or42a1* in the adult zebrafish spinal cord, specifically within Sox2^+^ ERG progenitor cells at the ependymal canal, extending the emergent paradigm of non-canonical OR expression to the injured central nervous system.

The functional link between *or42a1* expression and ERG progenitor cell cycle progression demonstrated through GO enrichment analysis, siRNA-mediated knockdown, CyclinD1-based G1/S transition assessment, PCNA-based proliferation quantification, and *or42a1* CDS mimic rescue experiments—firmly establishes *or42a1* as a positive regulator of NPC proliferation in the regenerating spinal cord. The reciprocal expression dynamics of dre-miR-N1 and *or42a1*, their direct molecular interaction validated by RNA EMSA and luciferase reporter assays, and the rescue of dre-miR-N1-mediated proliferative suppression upon *or42a1* CDS mimic co-delivery collectively delineate a coherent, functionally validated regulatory axis. This axis bears conceptual similarity to miRNA-mediated regulation of progenitor proliferation in other zebrafish regeneration paradigms: dre-miR-let7a suppresses Müller glia dedifferentiation by repressing ascl1a during retinal regeneration, and downregulation of miR-203 is required for Müller glia-derived progenitor proliferation in the same context (Rajaram et al., 2014; Ramachandran, Fausett and Goldman, 2015). In embryonic stem cells, miRNA-mediated post-transcriptional regulation actively promotes G1/S transition for rapid proliferative expansion (Wang et al., 2008), while hsa-miR-33 induces G1 arrest via CyclinD1 suppression during hepatic regeneration(Cirera-Salinas et al., 2012). Our findings extend this framework to spinal cord regeneration, demonstrating that progressive dre-miR-N1 downregulation in ventral ERG progenitor’s functions as a permissive signal for *or42a1* derepression, which in turn drives G1/S transition and proliferative expansion, a cascade essential for glial-bridge formation and functional locomotor recovery.

The mechanistic basis by which *or42a1*, a canonical GPCR, promotes ERG progenitor cell cycle entry remains to be fully resolved. Non-canonical OR signaling in non-olfactory tissues has been reported to engage cAMP-PKA, MAPK/ERK, and PI3K pathways (Oh, 2018), all of which intersect with core cell cycle regulatory machinery. Future studies examining the intracellular signaling cascades downstream of *or42a1* activation in ERG progenitors will be essential for fully resolving this regulatory axis at the molecular level. Additionally, while *ndr1* and *tshz1* exhibited weaker binding to dre-miR-N1 and expression dynamics inconsistent with direct miRNA regulation in the current experimental context, their upregulation during regeneration warrants further investigation as potential secondary targets shaping the broader regenerative transcriptional program.

## 4. Conclusion

This study delineates a previously uncharacterized miRNA-mediated regulatory axis governing ERG progenitor proliferation during spinal cord regeneration in adult zebrafish. Through comprehensive small RNA transcriptomics, bioinformatic target identification, and multiscale functional validation, we identify dre-miR-N1 as a novel, stage-specifically expressed miRNA that functions as a molecular brake on ERG progenitor proliferative activation. Its primary effector, *or42a1*, is an odorant receptor gene exhibiting ectopic, injury-inducible expression in Sox2^+^ ERG progenitor cells, the first reported instance of OR expression in the context of spinal cord regeneration. The reciprocal spatiotemporal expression dynamics of dre-miR-N1 and *or42a1*, validated by direct molecular interaction assays and rescue experiments, establish a dre-miR-N1/*or42a1* regulatory axis that coordinates progenitor cell cycle entry with the temporal demands of spinal cord regeneration.

The topology of this regulatory axis wherein progressive miRNA downregulation in an anatomically restricted progenitor compartment licenses target gene derepression to initiate proliferation suggests a conserved mechanism by which regenerative vertebrates temporally gate progenitor activation following CNS injury. The spatial precision of this regulation, with selective dre-miR-N1 attenuation in ventral ERGs undergoing EMT and proliferation while other ependymal compartments retain expression, implies that this miRNA–target interaction functions as a spatiotemporal checkpoint ensuring synchronized progenitor expansion calibrated to regenerative demand, thereby averting dysregulated proliferation that might impede functional recovery.

Beyond its implications for zebrafish regeneration biology, this study raises the intriguing possibility that modulating the miR-153 family, the closest annotated homolog of dre-miR-N1 or promoting non-canonical OR-mediated signaling in neural progenitors may represent viable therapeutic strategies for enhancing progenitor proliferation and functional recovery in mammalian SCI. Future investigations should elucidate the intracellular signaling cascades downstream of *or42a1* activation in ERG progenitors, define the upstream regulatory mechanisms governing dre-miR-N1 expression during regeneration, and determine whether analogous ectopic OR–miRNA regulatory interactions operate in mammalian CNS injury contexts. Such studies will be pivotal in translating the molecular insights derived from the zebrafish regeneration paradigm into clinically actionable therapeutic frameworks for the repair of spinal cord injury in future.

## 5. Materials and Methods

### 5.1. Experimental model and subject details

Adult AB-strain zebrafish (4-6 months old) of approximately equal sex ratio were used in this study which were maintained at a density of about 5 fish per litre water at 28°C in a custom aquarium, fed three times daily with daily water exchange. All husbandry and experiments were complied with institutional and national animal ethics guidelines and approved by the University Committee for the Purpose of Control and Supervision of Experiments on Animals (CPCSEA).

### 5.2. Injury procedures

Adult fish were anesthetized in 0.02% Tricaine methanesulfonate (MS222,Sigma Aldrich, Cat#A5040) and transection injury was performed as described previously (Hui and Ghosh, 2016). Briefly, a lateral incision was made on the skin at the level of the dorsal fin to expose the vertebral column and transection was made by using a micro scissor in the spinal cord. The injury-inflicted fish were transferred back to the housing system after the closure of the wound using Histoacryl gel (B. Braun, USA). Sham injury was carried out in order to analyse swimming behaviour as described here without spinal cord transection.

### 5.3. Total RNA isolation and mRNA-sequencing

The total RNA was extracted from zebrafish spinal cord at different time points post-injury using the TRIzol reagent in three biological replicates with spinal cord tissues pooled from 30 fish per replicate of each experimental time points. The RNA pools were normalized to 200 ng/µL concentration for every group and the quality was assessed using RNA HS assay kit (Thermofisher, Cat#Q32855) in a Qubit 4.0 fluorometer (Thermofisher, Cat#Q33238) following manufacturer’s protocol. Following qualification, it was used for library preparation and machine sequencing (IlluminaHSeq™ 2500 sequencing).

### 5.4. Small RNA library construction and sequencing

Small RNA Libraries from each experimental time points were prepared using NEBNext® Multiplex Small RNA Library Preparation kit (NEB, Cat#E7300L). Library quality was assessed by using DNA HS assay kit (Thermofisher, Cat#Q32851) in a Qubit 4.0 fluorometer as per the manufacturer’s protocol and all the libraries were normalized to 3 ng/µl and processed further for sequencing.

Small RNA sequencing data were analysed using the best-practice pipeline nf-core/smrnaseq with forward reads only (Ewels et al., 2020). Raw reads were assessed with FastQC v0.11.9 (Andrews, 2017),trimmed for low quality bases and adapter contaminations using fastp v.0.23.2 with default parameters and re-evaluated using FastQC (Chen et al., 2018). The genome fasta file and bowtie indexed files for Danio rerio (Source: NCBI, GCF_000002035.6_GRCz11), fasta files of miRBase mature and hairpin miRNA were obtained from AWS igenome (https://ewels.github.io/AWS-iGenomes/) repository. Adapter trimmed reads were mapped to Danio rerio miRBase mature hairpin miRNA sequences using Bowtie v1.3.1 (Langmead, 2010), and the resulting BAM files were sorted and indexed with SAMtools v1.14 (Li et al., 2009); alignment statistics were generated using SAMtools flagstat (Li et al., 2009). Known and novel miRNAs were identified using MiRDeep2 v2.0.1.2 (Friedländer et al., 2012), and differential expression was determined from absolute miR counts reported by smrnaseq pipeline using edgeR with exact Test (dispersion = 0.1) with miRNAs filtered at adjusted p-value <= 0.05 and Log2FC ± 2 (Anders et al., 2013; Chen et al., 2025).

### 5.5. Identification of target novel miRNAs

Raw sequencing reads were filtered to remove adaptor dimers, junk, low complexity, common RNA families and repeats, and only unique reads of 18–26 nucleotide (nt). These reads were mapped to zebrafish miRNA precursors miRBase 22.0 database (Kozomara, Birgaoanu and Griffiths-Jones, 2019), while unmapped reads were aligned using BLASTn to zebrafish genome (version CRCz11) and annotated against miRbase to identify candidate novel miRNAs. Novel candidates were further evaluated by miRDeep2, which models miRNA biogenesis by assessing read alignment to Dicer-processed hairpin structures, and providing log-odds scores to estimate the probability of the hairpin structures being genuine miRNA precursors. The analysis yielded a scored list of novel miRNAs with expression levels that met the stringent miRDeep2 criteria: cut-off score >4, significant randfold P value, and mature read count >10. Novel miRNAs identified at each time point were further compared using GeneVenn (http://genevenn.sourceforge.net/) to determine the target novel miRNAs that were common to all experimental time points, thereby reducing false positives. Among the novel miRNAs shared across all experimental groups, secondary hairpin structured were predicted using RNAfold module (Lorenz, Bernhart and Siederdissen, 2011), and the most stable stem loop conformations were defined as candidate novel miRNAs. Following the criteria by Zayed et al (Zayed, Qi and Peng, 2019), candidate pre-miRNAs were required to satisfy the following structural thresholds :(1) bulge size in the stem ≤ 12; (2) the stem region base-pairing ≥16; (3) minimum free energy (kcal/mol) ≤ -15; (4) hairpin length including all stem region and terminal loop ≥50 nt; (5) length of hairpin loop ≤ 20; (6) bulge size from the mature region ≤ 8; (7) biased error in a single bulge ≤ 4; (8) biased bulges in mature region ≤ 2; (9) total errors in the mature region ≤ 7; (10) mature sequence base-pairing ≥12; and (11) mature stem loop sequence occupancy ≥80.

### 5.6. miRNA seed sequence family analysis

For the assessment whether the candidate novel miRNA belongs to any existing annotated miRNA family irrespective of any species specificity, the mature sequence of the candidate novel miRNA was tested by using miRBase BLASTn function (Kozomara and Griffiths-Jones, 2011). The output of the search is the E-value, indicative of true sequence homology rather than any chance. Any result with an E-value <1 is regarded as a genuine hit for true sequence homology. The E-value threshold in this study was set to 10, for identification of any low-confidence similarity which may show any conserved seed sequence similarity to the input candidate novel miRNA (Szakats, McAtamney and Wilson, 2024).

### 5.7. In silico Target prediction and functional annotation of miRNAs

Two bioinformatics software tools, TargetScan and DIANA microT-CDS v. 5.0 were utilized to predict the possible target genes specific for the miRNA with minimum seed region homology to the target novel miRNA by applying following criteria :1) the miRNA sequence; 2) the 3′-UTR sequence of the database genes;3) the Watson–Crick base pairing in 5′ seed region of the target miRNA;4) minimum free energy required for binding;5) evolutionary conservation of the interaction . Target predictions with a threshold of context score < −0.2 for Targetscan and miTG score > 0.7 for DIANA micro T-CDS, respectively, were considered for further analysis (Disner et al., 2021). Prediction outputs from both the tools were then compared and the overlapping set of mutually predicted target genes were identified for the target miRNA.

### 5.8. *dicer*-Inhibitor application

1mg of Cib3b (HY-147918, MedChemExpress LLC, NJ, USA) was dissolved in 40 µL of 0.1% dimethylsulfoxide (DMSO), and soaked onto a small section of Gelfoam (Pfizer Inc., NY, USA) at a volume of 30 μl each. This Gelfoam piece was then divided into 30 smaller sections to yield dosage quantities ranging from 1 μl per piece, which were subsequently allowed to dry. One piece of the pre-soaked Cib3b was then placed at the site of injury after transection in each fish followed by sealing the wound using Histoacryl gel (B. Braun, USA). The gel-foam inserted fish were then reintroduced in the aquatic condition for survival and this protocol was systematically repeated over the following 6 days (up to 7 dpi). A similar procedure was employed for the control group of fish by administration of gel-foam soaked with 0.1% DMSO as a vehicle.

### 5.9. Quantification of miRNA by Universal Stem-loop qPCR

A Universal Stem-loop qRT-PCR was performed to quantify the relative miRNA expression levels in zebrafish tissue following a previously established protocol (Yang et al., 2014). Small RNAs were isolated from zebrafish tissue using mirVana™ miRNA Isolation Kit (ThermoFisher Scientific) in accordance with the manufacturer’s protocol and after confirming the RNA integrity using nanodrop spectrophotometer (Agilent Technologies) was transcribed into cDNA by using a universal stem-loop primer (USLP) which was designed from rice genome to avoid non-specific amplification as suggested by Yang et al (Yang et al., 2014). The miRNA-specific stem-loop RT primers were designed with the software miRPrimer2 and the stem-loop qRTPCR was performed in a LightCycler 480 system (Roche, Basel, Switzerland) using SYBR^TM^ Power Master Mix (Applied Biosystems, CA, USA) under standard cycling condition. Relative miRNA expression levels were quantified based on the cycle quantification (Cq) values and normalized to the reference gene U6 snRNA biogenesis 1 (*usb1*). Gene expression levels will be calculated using the 2^−ΔΔCq^ method. For target genes, beta-actin 2 (*actb2*) were employed as reference gene for normalization.

### 5.10. miRNA mimic synthesis and application

Target novel miRNA-specific mimic was synthesized using in vitro transcription method as per the previously established protocol (Sata et al., 2025). Firstly, the in vitro transcription templates for the mimic specific to the novel miRNA was designed using Primer3 software (Untergasser et al., 2012) and procured commercially. These templates were utilized for the production of miRNA guide and passenger strands by in vitro transcription method using mMESSAGE mMACHINE T3 IVT kit (ThermoFisher, Cat#AM1348). These mimic guide and passenger strands were purified using silica columns and annealed to produce functional mimic to the target novel miRNA.

Intraspinal delivery of novel miRNA mimic was performed in adult zebrafish using a modified protocol (Fan et al., 2024). Briefly, 2 µg of miR-N1 mimic was mixed with a transfection agent, Brainfectin (OZ Biosciences, Cat#IV-BF30100) to a final concentration of 20 pmol per 15 µl solution, loaded onto Gelfoam pieces (Pfizer Inc., NY, USA) at a volume of 30 μl each and subsequently allowed to dry followed division of the piece into 15-30 smaller sections which can hold mimic solution ranging from 1-2 μl per piece. The insertion of the Gelfoam was as described as in case of Cib3b treatment up to 7 dpi. Scrambled mimic was used as negative control and applied through the same process in control groups.

### 5.11. *or42a1* mRNA mimic synthesis and application

The CDS of *or42a1* transcript lacking the target miRNA binding site was synthesized by in vitro transcription method as described by a previously described protocol (Sata et al., 2025). Intraspinal application of *or42a1* mRNA mimic was performed as described previously in case of miRNA mimic application by gel foam method.

### 5.12. RNA Electrophoretic Mobility Shift Assays (EMSA)

Both the mimic sequence of the target miR-N1 and the 3’UTR regions of the target genes containing the binding site complementary to the seed region of the novel miRNA were synthesized by in vitro transcription method. Prior to hybridization, both the miRNA and mRNA oligonucleotides were denatured by heating at 80 °C for 10 min and subsequent incubation on ice to prevent secondary structure formation. For each 10 µl binding reaction, 200 nM miRNA oligonucleotide were incubated with varying concentrations of mRNA oligonucleotide (0 µM, 5 µM,10 µM and 20 µM, respectively) in EMSA binding buffer at 25 °C for 25 min. The resulting complexes were separated on an agarose gel by electrophoresis at 4 °C.

### 5.13. Dual-Luciferase Reporter Gene Assay

According to the *in silico* target prediction, 3’UTR sequence of the target gene containing the dre-miR-N1 binding site was amplified from the 5 dpf zebrafish spinal cord using the PCR method and cloned downstream of the Firefly luciferase coding sequence in the psicheck-2^TM^ vector (Promega). A corresponding mutant construct carrying a point-mutation within the seed-matching region was synthesized by site-directed mutagenesis commercially.

For in vitro assay, HEK293T cells were cultured following standard protocol (Mosmann, 1983) and co-transfected with the reporter plasmid (Wildtype or mutant) and either miRNA mimic or scrambled mimic for negative control using Lipofectamine 3000^TM^ (Thermo Fisher) according to the manufacturer’s protocol. After 24-48 hours post-transfection, luciferase activity was measured using Dual Glo Luciferase Assay System (Promega) in Varioskan LUX multimode microplate reader (Thermo Fisher scientific, Waltham, MA, USA) as per manufacturer’s instructions.

### 5.14. Target gene specific In Situ Hybridization and LNA probe mediated Fluorescent in situ hybridization for target miRNA

In situ hybridization using Digoxigenin (DIG)-labelled riboprobes was performed on spinal cord tissues fixed with 4% paraformaldehyde for 24 hours at 4°C and cryosection at 10 µm after embedding in tissue freezing medium. Tissue sections were washed twice with PBS for 5 min followed by target retrieval in citrate buffer for 5 min at 98°C. After retrieval, tissue sections were digested with Proteinase K solution (10 µg/ml) for permeabilization followed by hybridization of the sections with target gene-specific riboprobe diluted in hybridization buffer overnight at 70°C. After hybridization, sections were incubated with Anti-Digoxigenin antibody at 4°C for overnight followed by detection of signal using NBT-BCIP substrate. The *or42a1* specific DIG-labelled RNA probes used in this study were designed as per the standard protocol and synthesized using mMESSAGE mMACHINE T3 IVT kit (ThermoFisher, Cat#AM1348). The hybridized tissue sections were further processed for immunofluorescence staining using different antibodies to visualize the transcript expression in different cellular level as described previously (Lei et al., 2018).

For Fluorescent in situ hybridization (FISH) to detect miRNA expression in tissue sections, cryosections were hybridized after permeabilization using Fluorescent labelled Locked nucleic acid (LNA) Probe diluted in hybridization buffer at 70°C overnight and counterstained with DAPI (Silahtaroglu et al., 2007). Locked nucleic acid (LNA) probe complementary to the target novel miRNA was designed as per the protocol (Kloosterman et al., 2006) and Cy5-labelled LNA probe complementary to the target novel miRNA was synthesized commercially.

### 5.15. Swim path tracking

Zebrafish from control and various experimental groups were shifted to an opaque glass tank (60 cm in length and 30 cm in width) filled with aquarium water of 10 cm depth for acclimatization up to 15-20 minutes followed by a 30-minute video recording of swimming of zebrafish was made using a video camera (Sony, DSC-W830).

### 5.16. Histology and immunohistochemistry

Spinal cord tissues from control various experimental conditions were dissected out and fixed in 4% paraformaldehyde (Sigma) for overnight at 4°C followed by cryosectioning to 10-20 µm after embedding in Tissue freezing medium (Leica).

Immunofluorescence staining was performed in paraformaldehyde fixed 10 μm cryosections as described (Hui et al., 2017). Details of primary and secondary antibodies used in immunofluorescence were mentioned in the Key Resource Table. Briefly, Cryosections were rehydrated and given several washes in PBS with 0.1% Tween-20 (PBST). Antigen retrieval was done by keeping the slides in 90°C water bath for 15 min in sodium-citrate buffer (pH 6.0) followed by incubation with blocking solution (5% horse serum, 1% BSA in PBST) for 1 hr at 37°C and then with primary antibody for overnight at 4°C. This was followed by washes in PBST and incubated with secondary antibody for 2 hr at room temperature and nuclei were counter-stained with DAPI (Sigma). Immunostained sections were mounted with Gelvatol mounting medium (Vector Labs, Burlingame, CA) after two washes in PBST and stored in dark for microscopy.

### 5.17. Microscopy

Immunostained tissue sections and FISH tissue sections were photographed by using an OLYMPUS BX53F2 fluorescent microscope (Olympus, Tokyo, Japan). Confocal images were taken with a Zeiss LSM 710 confocal microscope (Carl Zeiss AG, Germany). Tissue sections after in situ hybridization were also photographed using OLYMPUS BX53F2 microscope under brightfield settings.

### 5.18. Quantification and Statistical analysis

#### 5.18.1. Cell Number Quantification

Cells positive with different markers (TUNEL^+^, CyclinD1^+^, Sox2^+^, PCNA^+^) in regenerating spinal cord were imaged within the injured areas of the spinal cord (1328 x 2376 pixels) under 20x magnification and counted manually using ImageJ software (US National Institutes of Health, Bethesda, MD, USA). The number of cells positive with different markers was calculated by averaging the value counted from six sections for each spinal cord from different experimental condition.

#### 5.18.2. Fluorescent intensity measurement

Fluorescent intensity of Cy5-labelled LNA probe signals was measured by calculating the mean grey value from a specific region of interest (ROI, 150000 mm^2^) in 5-6 tissue sections for each spinal cord from different groups and the data was represented by averaging the value. The threshold was normalized for each section before measurement.

#### 5.18.3. Glial bridging measurement

Glial bridging was quantified as the ratio of the cross-sectional area at the injury site to that of the intact spinal cord (750 µm rostral to the injury site using ImageJ software, as described previously (Klatt Shaw et al., 2021) . For each experimental group, six representative sections were analyzed, and the resulting ratios were averaged to determine the percentage of glial bridging during spinal cord regeneration.

#### 5.18.4. Swim Path Measurement

Trajectories of the swimming movement were obtained from the video recording of swimming zebrafish using idTracker software (de Polavieja lab, Cajal Institute, Madrid, Spain). Total travelled distance (cm/30 min), Mean Velocity (cm/s), Mean Swim speed (cm/s) and Burst Frequency (Hz) were calculated from the trajectory values using open-source R-packages.

#### 5.18.5. Luciferase Activity Measurement

For all experimental groups, firefly luciferase activity was normalized to Renilla luciferase activity and Relative luciferase activity was expressed as a ratio of firefly to Renilla signals. All experiments were performed with five independent biological replicates.

#### 5.18.6. Statistical Analysis

All statistical values are represented as mean ± SEM. To determine the statistical significance, the P values were calculated either with Mann–Whitney U tests, Student’s t-tests, or Fisher’s exact tests and P values less than 0.05 considered as statistically significant. All qRT-PCR results were obtained from averaging the data from 3-5 biological replicates. Respective Fig. legends also showed the statistical methods including sample size (N) and P values.

## Author Contributions

SG and SPH developed concept and designed experiments. AB designed and supervised all the qRTPCR experiments. SKJ and SM designed and supervised all the cell culture experiments. SG conducted experiments, collected data, and performed analysis. SG and SPH wrote manuscript. SPH supervised all aspects of the project. All authors read and approved the final manuscript.

## Acknowledgments

We thank Department of Science and Technology-Science & Engineering Research Board (DST-SERB), Govt. of India for their financial assistance through the Core Research Grant (CRG/2022/001663) to SPH. SG is a recipient of the Senior Research Fellowship (SRF) award [09/028(1145)-2020-EMR-I] from the Council of Scientific & Industrial Research (CSIR), Govt. of India. The funder played no role in study design, data collection, analysis and interpretation of data, or the writing of this manuscript.

## Competing interests

All authors declare no financial or non-financial competing interests.

## Data availability

All data generated or analysed during this study are available from the corresponding author upon reasonable request.

## Ethical Approval

The study was approved by the animal ethics committee (No. IAEC-04/BIOCHEM/CU/02-2022/03-SH) of Department of Biochemistry, University of Calcutta (Calcutta, India).

---

**Supplementary Figure S1:**
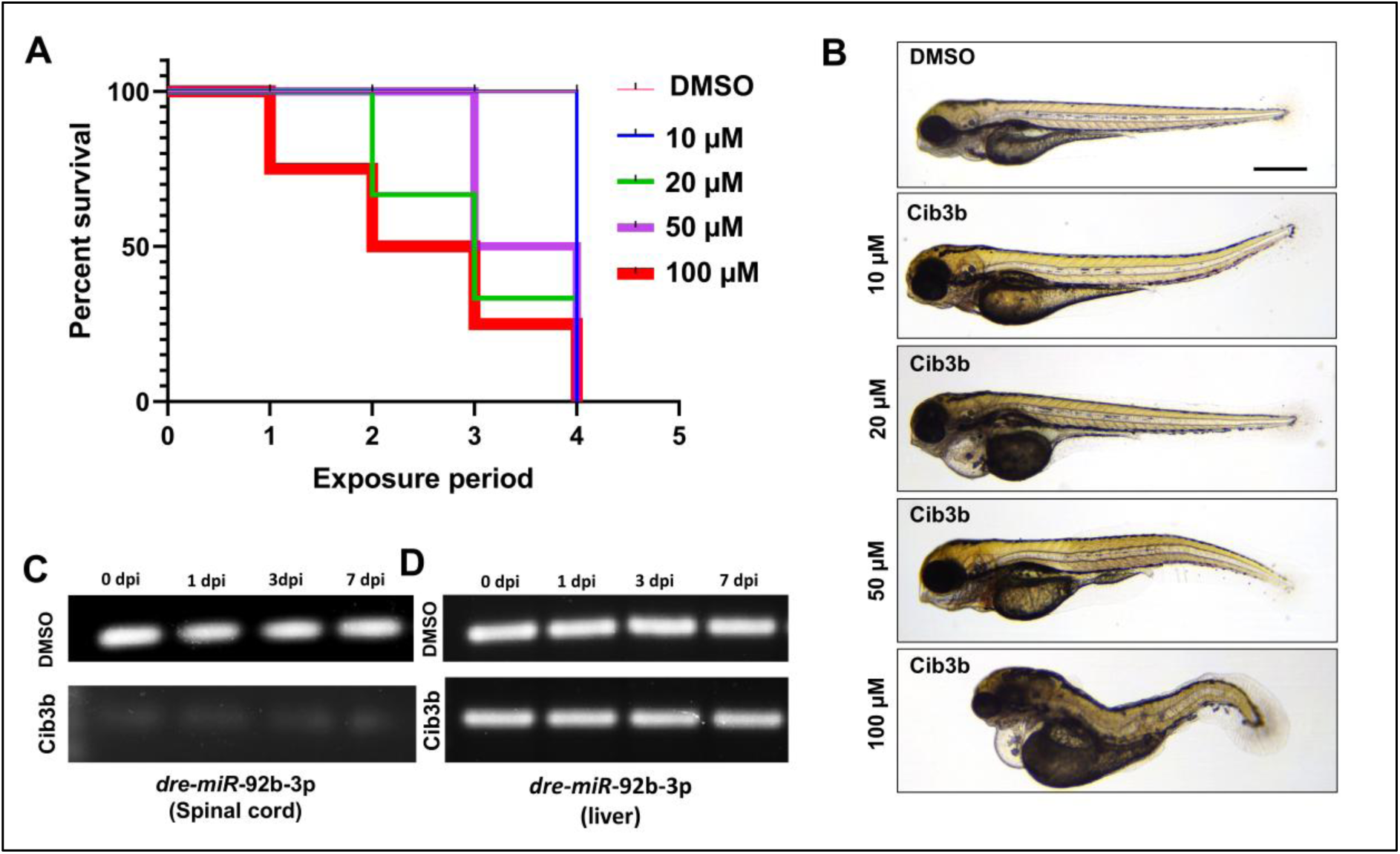
Dicer-inhibition supresses miRNA formation in zebrafish. (A) Kaplan–Meier survival curve of zebrafish embryos exposed to DMSO (controls) and 10µM, 20µM, 50µM and 100µM of Cib3b, during 4 days of treatment. Data are expressed as mean ± SE and n = 10 animals per group. (B) Morphological characteristics in control (DMSO-treated) and Cib3b-treated embryos. Anomalies in morphology of 96 hpf embryos were observed with increasing concentrations of Cib3b in experimental groups (N=10). (C) Gel images of transcript expression of a house-keeping miRNA, dre-miR-let7a, in uninjured and regenerating (1 dpi,3 dpi, and 7 dpi) spinal cord of adult zebrafish treated with DMSO and Cib3b. (D) Gel images of transcript expression of dre-miR-let7a, in liver tissue adjacent to the uninjured and regenerating (1 dpi,3 dpi, and 7 dpi) spinal cord of adult zebrafish treated with DMSO and Cib3b.

**Supplementary Figure S2:**
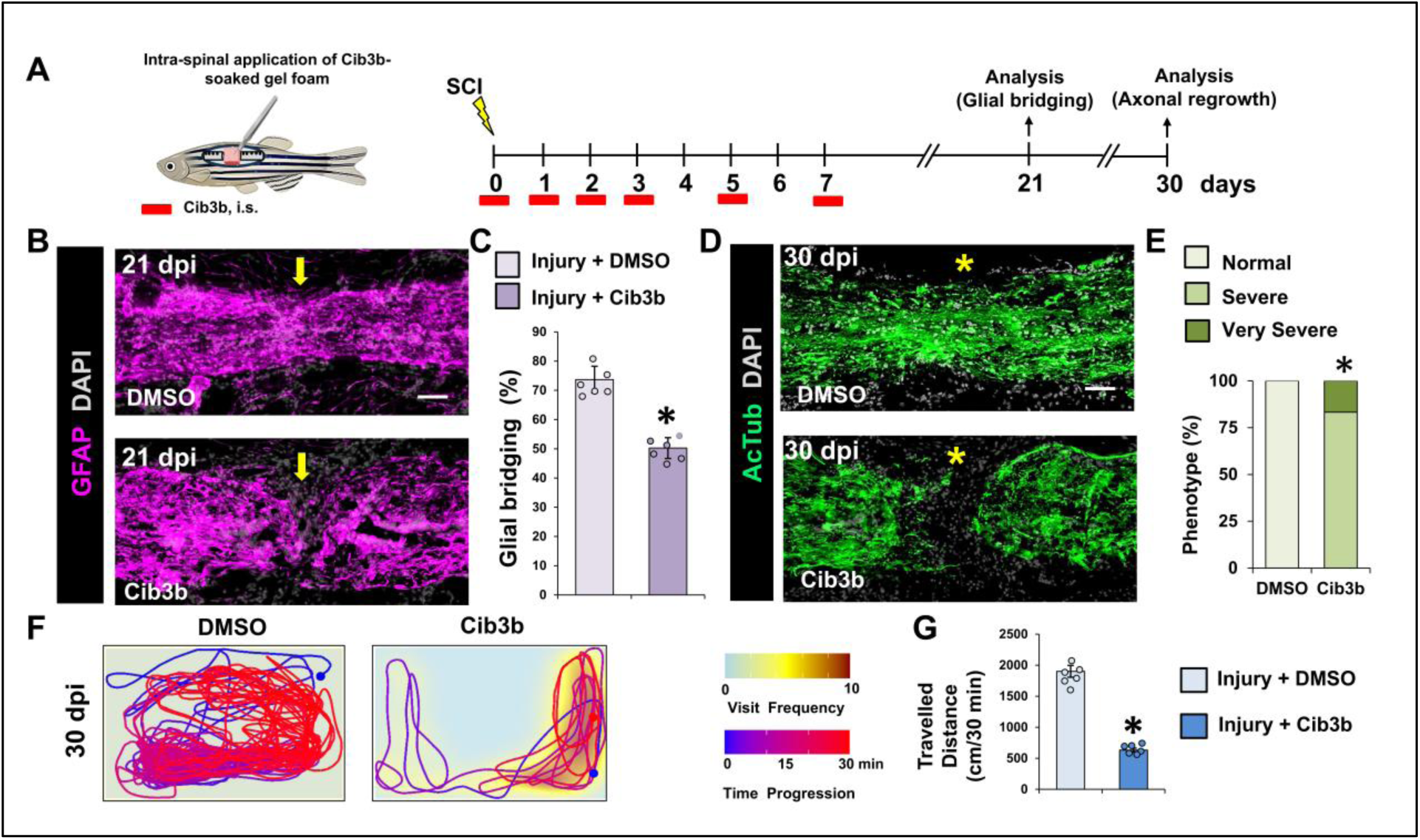
Suppression of miRNA formation by Dicer-inhibition impedes functional recovery after spinal cord injury in zebrafish. (A) Experimental intervention involving Cib3b treatment. i.s. intra-spinal injection. (B) Immunohistochemical staining of spinal cord sections after 21 dpi from DMSO-treated and Cib3b-treated fish showing glial bridge formation. Yellow arrow marked the glial bridging. (C)Quantification of glial bridging in (B) (N=6). (D) Immunohistochemical staining of spinal cord sections after 30 dpi from DMSO-treated (Upper) and Cib3b-treated (Lower) fish showing axonal projections. (E) Quantification of regeneration in (D) (N=5). (F) Swim tracking of individual animals after 30 dpi from DMSO-treated and Cib3b-treated experimental groups (N=8). (G) Quantification of total distance moved in (F). Yellow arrow indicates glial bridging; Asterisks indicate injury epicenter. Confocal projections of z stacks are shown. *p < 0.05; **p < 0.01; Fisher’s exact test (E) and Mann-Whitney U test (C, G). Magnification,10x; Scale bars, 50 µm (B, D) and 0.1 m (F). Yellow arrow indicates glial bridging; Asterisks indicate injury epicenter. Confocal projections of z stacks are shown. *p < 0.05; **p < 0.01; Fisher’s exact test (e) and Mann-Whitney U test (c, g). Magnification,10x; Scale bars, 50 µm (b, d) and 0.1 m (f).

**Supplementary Figure S3:**
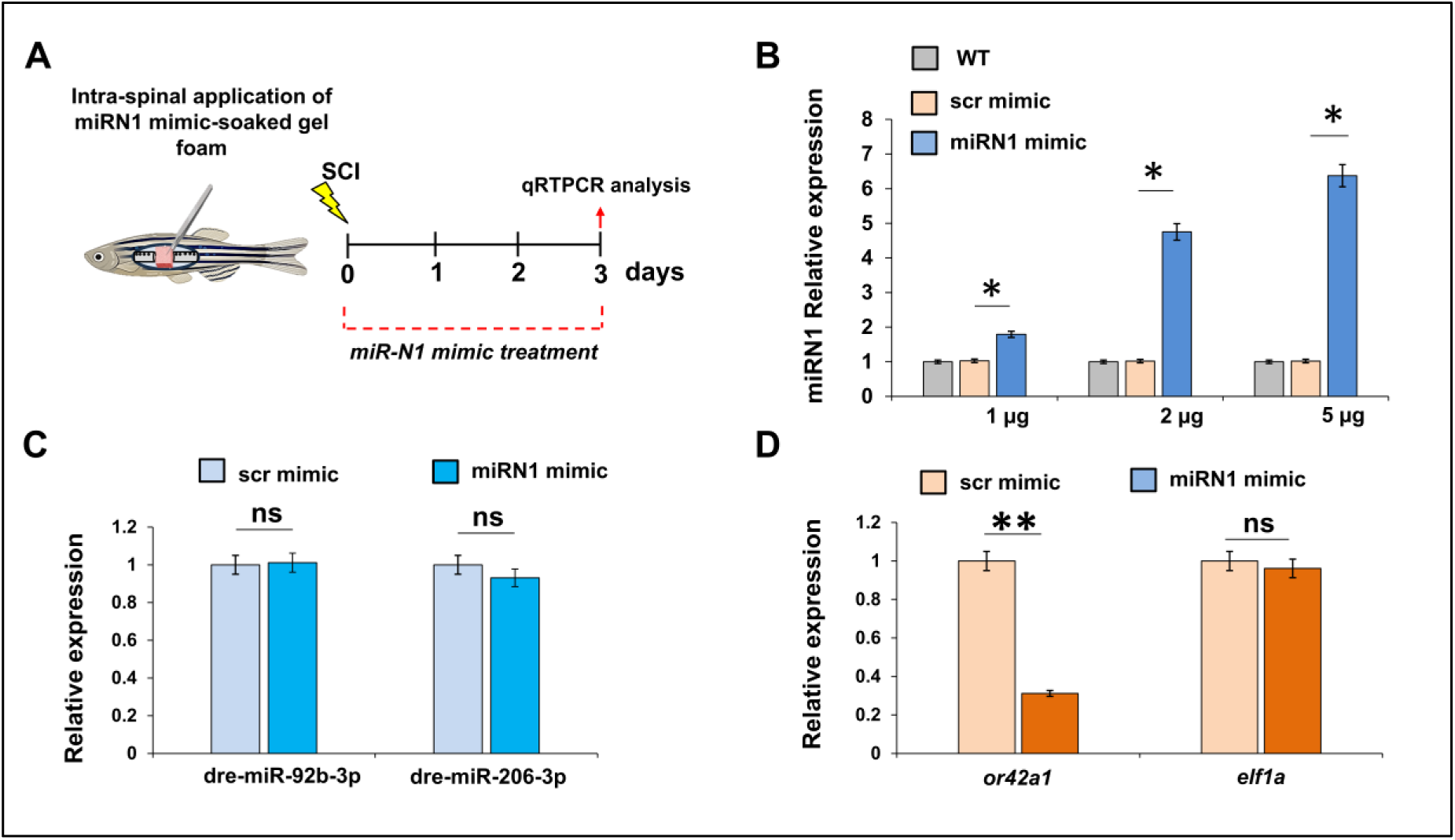
Dose dependent overexpression of miRN1 by specific mimic application. (A) Experimental intervention involving miRN1 mimic treatment by intra-spinal injection. (B) Quantification of dre-miR-N1 level after application of different doses (1 µg, 2 µg and 5 µg) of miRN1 specific mimic; (C) Quantification of the transcript level of two housekeeping miRNAs after miRN1 mimic treatment in comparison to scrambled-mimic treated group. (D) Relative expression analysis of *or42a1* and *eef1a1l1* after miRN1 mimic treated condition compared to scrambled treated control.

**Supplementary Figure S4:**
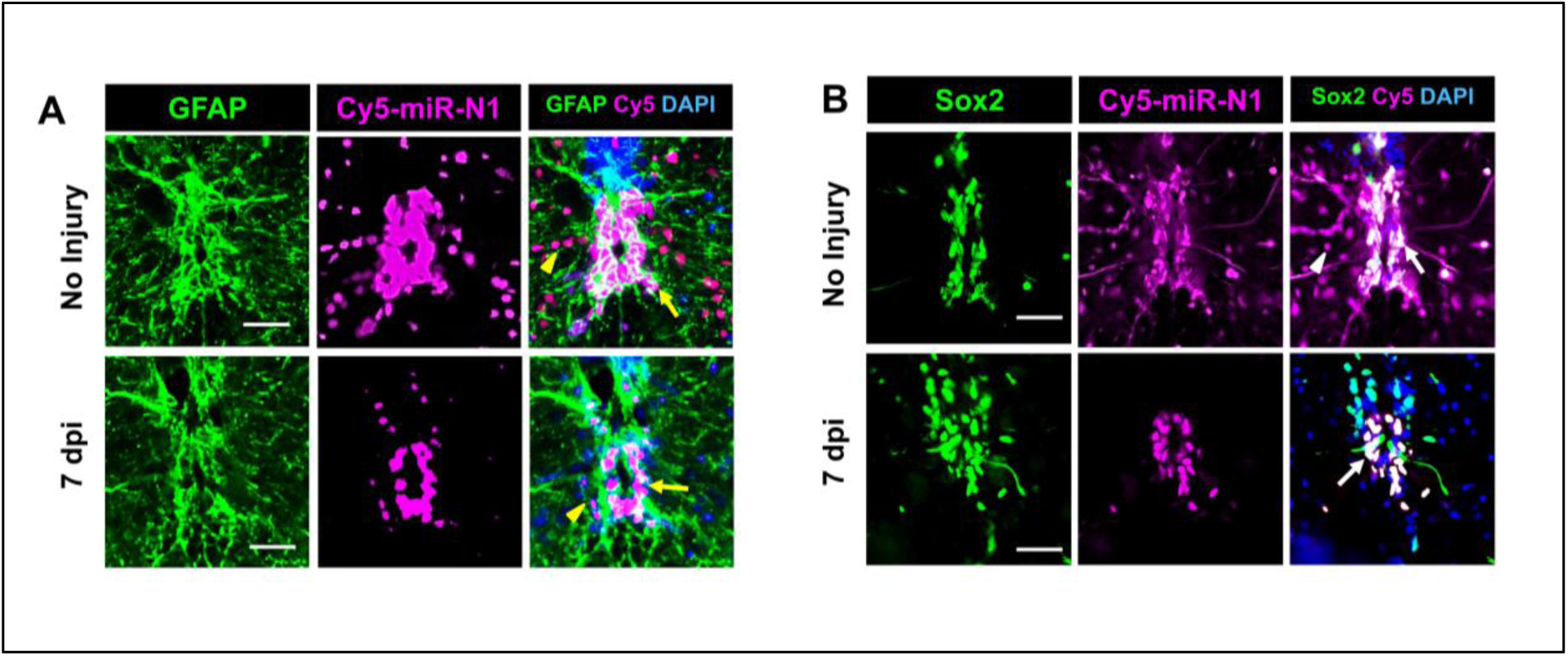
dre-miR-N1 colocalizes with ERG cells in spinal cord. Immunohistochemical staining of spinal cord (transverse Section) in uninjured and 7 dpi experimental groups showing (A) expression of GFAP, Cy5-labelled dre-miR-N1 and their colocalization including DAPI. Yellow arrow marked the GFAP^+^ cells co-expressing dre-miR-N1; yellow arrow-head marked the dre-miR-N1 expression not colocalized with GFAP, and (B) expression of Sox2, Cy5-labelled dre-miR-N1 and their colocalization including DAPI. White arrow marked the Sox2^+^ cells co-expressing dre-miR-N1; white arrowhead marked the dre-miR-N1 expression not colocalized with Sox2. Magnification, 20x; Scale bar, 50µm.

**Supplementary Figure S5:**
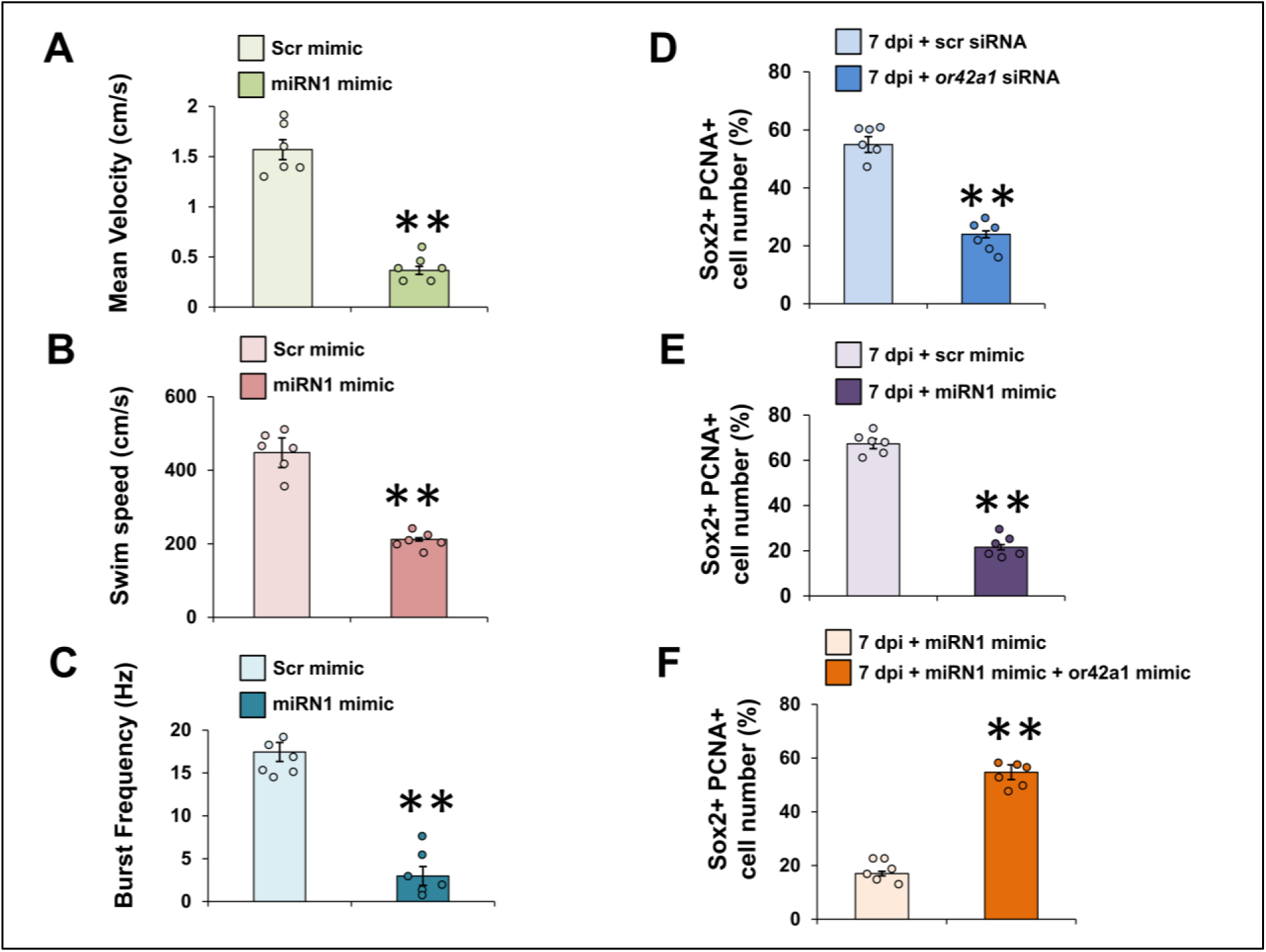
(A, B, C) Quantification of (A) Mean velocity (cm/s), (B) Swim speed (cm/s), and (C) Burst frequency (Hz) of scrambled mimic-treated and miRN1 mimic-treated fish. (D) Quantification of the number of Sox2^+^ PCNA^+^ cells after 7 dpi from scrambled siRNA-treated and *or42a1* siRNA-treated fish. (E) Quantification of the number of Sox2^+^ PCNA^+^ cells after 7 dpi from scrambled mimic-treated and miRN1 mimic-treated fish. (F) Quantification of the number of Sox2^+^ PCNA^+^ cells after 7 dpi from miRN1 mimic-treated and combined miRN1 mimic and *or42a1* CDS mimic-treated fish. Data represented as Mean±SEM, n=6; **p < 0.01; Mann-Whitney U test.

## Notes

### Competing Interest Statement

The authors have declared no competing interest.

## REFERENCES

1. Ahuja, C. S., Wilson, J. R., Nori, S., Kotter, M. R. N., Druschel, C., Curt, A. and Fehlings, M. G. (2017) “Traumatic spinal cord injury.,” Nature reviews. Disease primers, 3, p. 17018. doi: 10.1038/nrdp.2017.18.

2. Anders, S., McCarthy, D. J., Chen, Y., Okoniewski, M., Smyth, G. K., Huber, W. and Robinson, M. D. (2013) “Count-based differential expression analysis of RNA sequencing data using R and Bioconductor.,” Nature Protocols, 8(9), pp. 1765–1786. doi: 10.1038/nprot.2013.099.

3. Andrews, S. (2017) “FastQC: a quality control tool for high throughput sequence data. 2010.”

4. Anjum, A., Yazid, M. D., Fauzi Daud, M., Idris, J., Ng, A. M. H., Selvi Naicker, A., Ismail, O. H. R., Athi Kumar, R. K. and Lokanathan, Y. (2020) “Spinal cord injury: pathophysiology, multimolecular interactions, and underlying recovery mechanisms.,” International Journal of Molecular Sciences, 21(20). doi: 10.3390/ijms21207533.

5. Argo, S., Weth, F. and Korsching, S. I. (2003) “Analysis of penetrance and expressivity during ontogenesis supports a stochastic choice of zebrafish odorant receptors from predetermined groups of receptor genes.,” The European Journal of Neuroscience, 17(4), pp. 833–843. doi: 10.1046/j.1460-9568.2003.02505.x.

6. Barth, A. L., Dugas, J. C. and Ngai, J. (1997) “Noncoordinate expression of odorant receptor genes tightly linked in the zebrafish genome.,” Neuron, 19(2), pp. 359–369. doi: 10.1016/s0896-6273(00)80945-9.

7. Becker, C. G. and Becker, T. (2008) “Adult zebrafish as a model for successful central nervous system regeneration.,” Restorative Neurology and Neuroscience, 26(2-3), pp. 71–80.

8. Borger, A., Stadlmayr, S., Haertinger, M., Semmler, L., Supper, P., Millesi, F. and Radtke, C. (2022) “How miRNAs Regulate Schwann Cells during Peripheral Nerve Regeneration-A Systemic Review.,” International Journal of Molecular Sciences, 23(7). doi: 10.3390/ijms23073440.

9. Chen, L., Chuang, M., Koorman, T., Boxem, M., Jin, Y. and Chisholm, A. D. (2015) “Axon injury triggers EFA-6 mediated destabilization of axonal microtubules via TACC and doublecortin like kinase.,” eLife, 4. doi: 10.7554/eLife.08695.

10. Chen, S., Zhou, Y., Chen, Y. and Gu, J. (2018) “fastp: an ultra-fast all-in-one FASTQ preprocessor.,” Bioinformatics, 34(17), pp. i884–i890. doi: 10.1093/bioinformatics/bty560.

11. Chen, Y., Chen, L., Lun, A. T. L., Baldoni, P. L. and Smyth, G. K. (2025) “edgeR v4: powerful differential analysis of sequencing data with expanded functionality and improved support for small counts and larger datasets.,” Nucleic Acids Research, 53(2). doi: 10.1093/nar/gkaf018.

12. Cirera-Salinas, D., Pauta, M., Allen, R. M., Salerno, A. G., Ramírez, C. M., Chamorro-Jorganes, A., Wanschel, A. C., Lasuncion, M. A., Morales-Ruiz, M., Suarez, Y., Baldan, Á., Esplugues, E. and Fernández-Hernando, C. (2012) “Mir-33 regulates cell proliferation and cell cycle progression.,” Cell Cycle, 11(5), pp. 922–933. doi: 10.4161/cc.11.5.19421.

13. Dhahbi, J. M., Atamna, H., Boffelli, D., Magis, W., Spindler, S. R. and Martin, D. I. K. (2011) “Deep sequencing reveals novel microRNAs and regulation of microRNA expression during cell senescence.,” Plos One, 6(5), p. e20509. doi: 10.1371/journal.pone.0020509.

14. Ding, L. Z., Xu, J., Yuan, C., Teng, X. and Wu, Q. M. (2020) “MiR-7a ameliorates spinal cord injury by inhibiting neuronal apoptosis and oxidative stress.,” European review for medical and pharmacological sciences, 24(1), pp. 11–17. doi: 10.26355/eurrev_202001_19890.

15. Disner, G. R., Falcão, M. A. P., Lima, C. and Lopes-Ferreira, M. (2021) “In Silico Target Prediction of Overexpressed microRNAs from LPS-Challenged Zebrafish (Danio rerio) Treated with the Novel Anti-Inflammatory Peptide TnP.,” International Journal of Molecular Sciences, 22(13). doi: 10.3390/ijms22137117.

16. Ewels, P. A., Peltzer, A., Fillinger, S., Patel, H., Alneberg, J., Wilm, A., Garcia, M. U., Di Tommaso, P. and Nahnsen, S. (2020) “The nf-core framework for community-curated bioinformatics pipelines.,” Nature Biotechnology, 38(3), pp. 276–278. doi: 10.1038/s41587-020-0439-x.

17. Fan, J., Liu, X., Duan, Z., Zhao, H., Chang, Z. and Li, L. (2024) “The Regulatory Role of miRNAs in Zebrafish Fin Regeneration.,” International Journal of Molecular Sciences, 25(19). doi: 10.3390/ijms251910542.

18. Franco, R., Garrigós, C., Capó, T., Serrano-Marín, J., Rivas-Santisteban, R. and Lillo, J. (2025) “Olfactory receptors in neural regeneration in the central nervous system.,” Neural regeneration research, 20(9), pp. 2480–2494. doi: 10.4103/NRR.NRR-D-24-00495.

19. Friedländer, M. R., Mackowiak, S. D., Li, N., Chen, W. and Rajewsky, N. (2012) “miRDeep2 accurately identifies known and hundreds of novel microRNA genes in seven animal clades.,” Nucleic Acids Research, 40(1), pp. 37–52. doi: 10.1093/nar/gkr688.

20. Galagali, H. (2023) “Regulation of microRNAs during development and environmental stress in Caenorhabditis elegans.”

21. Guo, S., Redenski, I. and Levenberg, S. (2021) “Spinal cord repair: from cells and tissue engineering to extracellular vesicles.,” Cells, 10(8). doi: 10.3390/cells10081872.

22. Gupta, S. and Hui, S. P. (2025) “Epigenetic Cross-Talk Between Sirt1 and Dnmt1 Promotes Axonal Regeneration After Spinal Cord Injury in Zebrafish.,” Molecular Neurobiology, 62(2), pp. 2396–2419. doi: 10.1007/s12035-024-04408-w.

23. Huang, R., Chen, M., Yang, L., Wagle, M., Guo, S. and Hu, B. (2017) “MicroRNA-133b Negatively Regulates Zebrafish Single Mauthner-Cell Axon Regeneration through Targeting tppp3 in Vivo.,” Frontiers in Molecular Neuroscience, 10, p. 375. doi: 10.3389/fnmol.2017.00375.

24. Hui, S. and Ghosh, S. (2016) “Various modes of spinal cord injury to study regeneration in adult zebrafish,” Bio-protocol, 6(23). doi: 10.21769/BioProtoc.2043.

25. Hui, S. P., Dutta, A. and Ghosh, S. (2010) “Cellular response after crush injury in adult zebrafish spinal cord.,” Developmental Dynamics, 239(11), pp. 2962–2979. doi: 10.1002/dvdy.22438.

26. Hui, S. P., Sheng, D. Z., Sugimoto, K., Gonzalez-Rajal, A., Nakagawa, S., Hesselson, D. and Kikuchi, K. (2017) “Zebrafish Regulatory T Cells Mediate Organ-Specific Regenerative Programs.,” Developmental Cell, 43(6), pp. 659–672.e5. doi: 10.1016/j.devcel.2017.11.010.

27. Jiang, D., Gong, F., Ge, X., Lv, C., Huang, C., Feng, S., Zhou, Z., Rong, Y., Wang, J., Ji, C., Chen, J., Zhao, W., Fan, J., Liu, W. and Cai, W. (2020) “Neuron-derived exosomes-transmitted miR-124-3p protect traumatically injured spinal cord by suppressing the activation of neurotoxic microglia and astrocytes.,” Journal of nanobiotechnology, 18(1), p. 105. doi: 10.1186/s12951-020-00665-8.

28. Jundi, D., Coutanceau, J.-P., Bullier, E., Imarraine, S., Fajloun, Z. and Hong, E. (2023) “Expression of olfactory receptor genes in non-olfactory tissues in the developing and adult zebrafish.,” Scientific Reports, 13(1), p. 4651. doi: 10.1038/s41598-023-30895-3.

29. Klatt Shaw, D., Saraswathy, V. M., Zhou, L., McAdow, A. R., Burris, B., Butka, E., Morris, S. A., Dietmann, S. and Mokalled, M. H. (2021) “Localized EMT reprograms glial progenitors to promote spinal cord repair.,” Developmental Cell, 56(5), pp. 613–626.e7. doi: 10.1016/j.devcel.2021.01.017.

30. Kloosterman, W. P., Wienholds, E., de Bruijn, E., Kauppinen, S. and Plasterk, R. H. A. (2006) “In situ detection of miRNAs in animal embryos using LNA-modified oligonucleotide probes.,” Nature Methods, 3(1), pp. 27–29. doi: 10.1038/nmeth843.

31. Kozomara, A., Birgaoanu, M. and Griffiths-Jones, S. (2019) “miRBase: from microRNA sequences to function.,” Nucleic Acids Research, 47(D1), pp. D155–D162. doi: 10.1093/nar/gky1141.

32. Kozomara, A. and Griffiths-Jones, S. (2011) “miRBase: integrating microRNA annotation and deep-sequencing data.,” Nucleic Acids Research, 39(Database issue), pp. D152–7. doi: 10.1093/nar/gkq1027.

33. Kuscha, V., Frazer, S. L., Dias, T. B., Hibi, M., Becker, T. and Becker, C. G. (2012) “Lesion-induced generation of interneuron cell types in specific dorsoventral domains in the spinal cord of adult zebrafish.,” The Journal of Comparative Neurology, 520(16), pp. 3604–3616. doi: 10.1002/cne.23115.

34. Langmead, B. (2010) “Aligning short sequencing reads with Bowtie.,” *Current Protocols in Bioinformatics*, Chapter 11, p. Unit 11.7. doi: 10.1002/0471250953.bi1107s32.

35. Lei, Z., van Mil, A., Xiao, J., Metz, C. H. G., van Eeuwijk, E. C. M., Doevendans, P. A. and Sluijter, J. P. G. (2018) “MMISH: Multicolor microRNA in situ hybridization for paraffin embedded samples.,” *Biotechnology reports (Amsterdam*, Netherlands), 18, p. e00255. doi: 10.1016/j.btre.2018.e00255.

36. Li, H., Handsaker, B., Wysoker, A., Fennell, T., Ruan, J., Homer, N., Marth, G., Abecasis, G., Durbin, R. and 1000 Genome Project Data Processing Subgroup (2009) “The Sequence Alignment/Map format and SAMtools.,” Bioinformatics, 25(16), pp. 2078–2079. doi: 10.1093/bioinformatics/btp352.

37. Li, P., Teng, Z.-Q. and Liu, C.-M. (2016) “Extrinsic and Intrinsic Regulation of Axon Regeneration by MicroRNAs after Spinal Cord Injury.,” Neural plasticity, 2016, p. 1279051. doi: 10.1155/2016/1279051.

38. Li, X., Gao, Y., Tian, F., Du, R., Yuan, Y., Li, P., Liu, F. and Wang, C. (2021) “miR-31 promotes neural stem cell proliferation and restores motor function after spinal cord injury.,” Experimental Biology and Medicine, 246(11), pp. 1274–1286. doi: 10.1177/1535370221997071.

39. Lin, S., Xu, C., Lin, J., Hu, H., Zhang, C. and Mei, X. (2021) “Regulation of inflammatory cytokines for spinal cord injury recovery.,” Histology and Histopathology, 36(2), pp. 137–142. doi: 10.14670/HH-18-262.

40. Liu, H., Xu, X., Tu, Y., Chen, K., Song, L., Zhai, J., Chen, S., Rong, L., Zhou, L., Wu, W., So, K.-F., Ramakrishna, S. and He, L. (2020) “Engineering Microenvironment for Endogenous Neural Regeneration after Spinal Cord Injury by Reassembling Extracellular Matrix.,” ACS Applied Materials & Interfaces, 12(15), pp. 17207–17219. doi: 10.1021/acsami.9b19638.

41. Lorenz, R., Bernhart, S. H. and Siederdissen, C. H. zu (2011) “ViennaRNA package 2.0. algorithms,” Mol Biol.

42. Lu, T. X. and Rothenberg, M. E. (2018) “MicroRNA.,” The Journal of Allergy and Clinical Immunology, 141(4), pp. 1202–1207. doi: 10.1016/j.jaci.2017.08.034.

43. McCreight, J. C., Schneider, S. E., Wilburn, D. B. and Swanson, W. J. (2017) “Evolution of microRNA in primates.,” Plos One, 12(6), p. e0176596. doi: 10.1371/journal.pone.0176596.

44. Milbreta, U., Lin, J., Pinese, C., Ong, W., Chin, J. S., Shirahama, H., Mi, R., Williams, A., Bechler, M. E., Wang, J., Ffrench-Constant, C., Hoke, A. and Chew, S. Y. (2019) “Scaffold-Mediated Sustained, Non-viral Delivery of miR-219/miR-338 Promotes CNS Remyelination.,” Molecular Therapy, 27(2), pp. 411–423. doi: 10.1016/j.ymthe.2018.11.016.

45. Mokalled, M. H., Patra, C., Dickson, A. L., Endo, T., Stainier, D. Y. R. and Poss, K. D. (2016) “Injury-induced ctgfa directs glial bridging and spinal cord regeneration in zebrafish.,” Science, 354(6312), pp. 630–634. doi: 10.1126/science.aaf2679.

46. Mosmann, T. (1983) “Rapid colorimetric assay for cellular growth and survival: application to proliferation and cytotoxicity assays.,” Journal of Immunological Methods, 65(1-2), pp. 55–63. doi: 10.1016/0022-1759(83)90303-4.

47. Oh, S. J. (2018) “System-Wide Expression and Function of Olfactory Receptors in Mammals.,” Genomics & informatics, 16(1), pp. 2–9. doi: 10.5808/GI.2018.16.1.2.

48. Patel, A. and Peralta-Yahya, P. (2023) “Olfactory receptors as an emerging chemical sensing scaffold.,” Biochemistry, 62(2), pp. 187–195. doi: 10.1021/acs.biochem.2c00486.

49. Rajaram, K., Harding, R. L., Hyde, D. R. and Patton, J. G. (2014) “miR-203 regulates progenitor cell proliferation during adult zebrafish retina regeneration.,” Developmental Biology, 392(2), pp. 393–403. doi: 10.1016/j.ydbio.2014.05.005.

50. Ramachandran, R., Fausett, B. V. and Goldman, D. (2015) “Ascl1a regulates Müller glia dedifferentiation and retinal regeneration through a Lin-28-dependent, let-7 microRNA signalling pathway.,” Nature Cell Biology, 17(4), p. 532. doi: 10.1038/ncb3144.

51. Rehmsmeier, M., Steffen, P., Hochsmann, M. and Giegerich, R. (2004) “Fast and effective prediction of microRNA/target duplexes.,” RNA (New York), 10(10), pp. 1507–1517. doi: 10.1261/rna.5248604.

52. Reimer, M. M., Kuscha, V., Wyatt, C., Sörensen, I., Frank, R. E., Knüwer, M., Becker, T. and Becker, C. G. (2009) “Sonic hedgehog is a polarized signal for motor neuron regeneration in adult zebrafish.,” The Journal of Neuroscience, 29(48), pp. 15073–15082. doi: 10.1523/JNEUROSCI.4748-09.2009.

53. Sata, T. N., Ismail, M., Sah, A. K. and Venugopal, S. K. (2025) “Production of Functional miRNA Mimics Using In Vitro Transcription.,” Current protocols, 5(6), p. e70163. doi: 10.1002/cpz1.70163.

54. Sato, Y., Miyasaka, N. and Yoshihara, Y. (2007) “Hierarchical regulation of odorant receptor gene choice and subsequent axonal projection of olfactory sensory neurons in zebrafish.,” The Journal of Neuroscience, 27(7), pp. 1606–1615. doi: 10.1523/JNEUROSCI.4218-06.2007.

55. Shen, W., Cai, J., Li, J., Li, W., Shi, P., Zhao, X. and Feng, S. (2024) “Regulation of micrornas after spinal cord injury in adult zebrafish.,” Journal of Molecular Neuroscience, 74(3), p. 66. doi: 10.1007/s12031-024-02242-2.

56. Silahtaroglu, A. N., Nolting, D., Dyrskjøt, L., Berezikov, E., Møller, M., Tommerup, N. and Kauppinen, S. (2007) “Detection of microRNAs in frozen tissue sections by fluorescence in situ hybridization using locked nucleic acid probes and tyramide signal amplification.,” Nature Protocols, 2(10), pp. 2520–2528. doi: 10.1038/nprot.2007.313.

57. Szakats, S., McAtamney, A. and Wilson, M. J. (2024) “Identification of novel microRNAs in the embryonic mouse brain using deep sequencing.,” Molecular and Cellular Biochemistry, 479(2), pp. 297–311. doi: 10.1007/s11010-023-04730-2.

58. Untergasser, A., Cutcutache, I., Koressaar, T., Ye, J., Faircloth, B. C., Remm, M. and Rozen, S. G. (2012) “Primer3--new capabilities and interfaces.,” Nucleic Acids Research, 40(15), p. e115. doi: 10.1093/nar/gks596.

59. Wang, F., Zhang, C., Zhang, Q., Li, J., Xue, Y. and He, X. (2023) “Lithium ameliorates spinal cord injury through endoplasmic reticulum stress-regulated autophagy and alleviated apoptosis through IRE1 and PERK/eIF2α …,” Journal of ….

60. Wang, Y., Baskerville, S., Shenoy, A., Babiarz, J. E., Baehner, L. and Blelloch, R. (2008) “Embryonic stem cell-specific microRNAs regulate the G1-S transition and promote rapid proliferation.,” Nature Genetics, 40(12), pp. 1478–1483. doi: 10.1038/ng.250.

61. Xu, Y., Zhou, J., Liu, C., Zhang, S., Gao, F., Guo, W., Sun, X., Zhang, C., Li, H., Rao, Z., Qiu, S., Zhu, Q., Liu, X., Guo, X., Shao, Z., Bai, Y., Zhang, X. and Quan, D. (2021) “Understanding the role of tissue-specific decellularized spinal cord matrix hydrogel for neural stem/progenitor cell microenvironment reconstruction and spinal cord injury.,” Biomaterials, 268, p. 120596. doi: 10.1016/j.biomaterials.2020.120596.

62. Yang, L., Wang, S., Tang, L., Liu, B., Ye, W., Wang, L., Wang, Z., Zhou, M. and Chen, B. (2014) “Universal stem-loop primer method for screening and quantification of microRNA.,” Plos One, 9(12), p. e115293. doi: 10.1371/journal.pone.0115293.

63. Yu, Y.-M., Gibbs, K. M., Davila, J., Campbell, N., Sung, S., Todorova, T. I., Otsuka, S., Sabaawy, H. E., Hart, R. P. and Schachner, M. (2011) “MicroRNA miR-133b is essential for functional recovery after spinal cord injury in adult zebrafish.,” The European Journal of Neuroscience, 33(9), pp. 1587–1597. doi: 10.1111/j.1460-9568.2011.07643.x.

64. Zayed, Y., Qi, X. and Peng, C. (2019) “Identification of novel micrornas and characterization of microrna expression profiles in zebrafish ovarian follicular cells.,” Frontiers in endocrinology, 10, p. 518. doi: 10.3389/fendo.2019.00518.

