## Supplementary Materials for "A Novel MicroRNA-Odorant Receptor Axis Governs Neural Progenitor Cell Proliferation in Zebrafish CNS"

### Supplementary Table Legends

**Supplementary Table S1 Legend:** Mature sequence and minimum free energy (in kcal/mol) of 12 filtered novel miRNAs using RNAfold module. Top 3 stable miRNAs were marked in red.

### Supplementary Table S1:

| Name | Mature sequence (5'-3') | MFE | miRdeep2 score |
|---|---|---|---|
| **miR-N1** | aucauuuuugugacuaugcaacu | -44.50 kcal/mol | 4186.4 |
| **miR-N2** | ucagcacucggacagccucuu | -27.50 kcal/mol | 2241 |
| **miR-N3** | uccaucagucacgugaccuacc | -30.90 kcal/mol | 2149.9 |
| **miR-N4** | ugcgcaugcgugaacuuuguacc | -1.93 kcal/mol | 2296.5 |
| **miR-N5** | uacacgagaacaccaggacaca | -17.70 kcal/mol | 1397.3 |
| **miR-N6** | uuuaaaaguguccuccagagc | -13.40 kcal/mol | 1322.1 |
| **miR-N7** | cuagcaccauuugaaaucgguc | -21.00 kcal/mol | 923.3 |
| **miR-N8** | agacucuccaguacacuggcc | -24.90 kcal/mol | 839.1 |
| **miR-N9** | ucaaaaguguaccgaaccaaac | -24.70 kcal/mol | 615.6 |
| **miR-N10** | uacgguuugguaagcuuuuaug | -23.60 kcal/mol | 486.2 |
| **miR-N11** | uuacaauuaaaggauauuucu | -17.00 kcal/mol | 425.1 |
| **miR-N12** | acuggcacucauugacuucugu | -17.00 kcal/mol | 345.5 |

### Key Resource Table

| Reagent/ Resource | Source | Identifier |
|---|---|---|
| **Experimental subjects** | | |
| AB strain zebrafish | | RRID: ZIRC_ZL1 |
| HEK293T cell | | RRID: CVCL_0063 |
| psiCHECK™-2 Vector | Promega | Cat#C8021 |
| **Antibodies** | | |
| Rabbit polyclonal anti-GFAP | Dako | RRID: AB_2811722 (Cat#GA524) |
| Mouse monoclonal anti-acetylated tubulin | Sigma-Aldrich | RRID: AB_477585 Cat#T6793 |
| Rabbit polyclonal anti-Cyclin D1 | AnaSpec | RRID:AB_10561867 Cat#55399 |
| Rabbit polyclonal anti-SOX2 | Abcam | RRID:AB_2341193 Cat#ab97959 |
| Mouse monoclonal anti-PCNA | Abcam | RRID: AB_303394 Cat#ab29 |
| Goat anti-rabbit Alexa Fluor 488, IgG (H+L) | Jackson Immunoresearch | RRID: AB_2338046 Cat#111-545-003 |
| Goat anti-mouse Alexa Fluor 488, IgG (H+L) | Jackson Immunoresearch | RRID: AB_2338845 Cat#115-545-062 |
| Goat anti-rabbit Alexa Fluor 594, IgG (H+L) | Jackson Immunoresearch | RRID: AB_2307325 Cat#111-585-144 |
| Goat anti-mouse Alexa Fluor 594, IgG (H+L) | Jackson Immunoresearch | RRID: AB_2338883 Cat#115-585-166 |
| Anti-Digoxigenin-AP, Fab fragments | Roche | RRID: AB_514497 Cat# 11093274910 |
| **Reagents** | | |
| Tissue Freezing Medium | Leica Biosystems | Cat#14020108926 |
| DAPI (4',6- diamidino-2-phenylindole dihydrochloride) | Sigma-Aldrich | Cat# D9542 |
| TRIzol Reagent | Thermo Fisher Scientific | Cat# 15596026 |
| Dimethyl Sulfoxide (DMSO) | Sigma-Aldrich | Cat# D4540 |
| Fetal Bovine Serum | Thermo Fisher Scientific | Cat# 10082139 |

| | | |
|---|---|---|
| Cib3b | MedchemExpress | Cat# HY-147918 |
| Histoacryl gel | B. Braun | Cat# 1050044 |
| Gelvatol mounting medium | Sigma-Aldrich | Cat# MW 4-88 |
| SIGMAFAST™ BCIP®/NBT Tablet | Sigma-Aldrich | Cat# B5655 |
| **Critical Commercial Assays** | | |
| GoScript™ Reverse Transcriptase Kit | Promega | Cat#A5003 |
| GoTaq Green Master Mix | Promega | Cat#M7122 |
| PrimeSTAR GXL DNA Polymerase | Takara Clonetech | Cat#R051A |
| SYBR™ Power Master Mix | Thermo Fisher Scientific | Cat# 4472908 |
| mMESSAGE mMACHINE™ T3 Transcription Kit | Thermo Fisher Scientific | Cat#AM1348 |
| RNA HS assay kit | Thermo Fisher Scientific | Cat# Q32855 |
| DNA HS assay kit | Thermo Fisher Scientific | Cat# Q32851 |
| NEBNext® Multiplex Small RNA Library Preparation kit | New England Biolab | Cat#E7300L |
| mirVana™ miRNA Isolation Kit | Thermo Fisher Scientific | Cat#AM1560 |
| In Situ Cell Death Detection Kit, Fluorescein | Roche | Cat# 11684795910 |
| Dual-Glo® Luciferase Assay System | Promega | Cat#E2920 |
| **Software and Algorithms** | | |
| FastQC V0.11.9 | (Andrews, 2017) | RRID:SCR_014583 |
| Fastp v0.23.2 | (Chen et al., 2018) | RRID:SCR_016962 |
| Bowtie v1.3.1 | (Langmead et al., 2009) | RRID:SCR_005476 |
| SAMtools v1.14 | (Li et al., 2009) | RRID:SCR_002105 |
| MiRDeep2 v2.0.1.2 | (Friedländer et al., 2012) | RRID:SCR_010829 |
| EdgeR v4.0 | (Chen et al., 2025) | RRID:SCR_012802 |
| RNAfold | (Lorenz, Bernhart and Siederdissen, 2011) | RRID:SCR_024427 |
| RNAhybrid | (Rehmsmeier et al., 2004) | RRID:SCR_003252 |
| miRBase | (Kozomara, Birgaoanu and Griffiths-Jones, 2019) | RRID:SCR_003152 |
| GeneVenn | | RRID:SCR_012117 |
| Primer3 | (Untergasser et al., 2012) | RRID:SCR_003139 |

| | | |
|---|---|---|
| idTracker | (Pérez-Escudero et al., 2014) | https://www.idtracker.es/ |
| Fiji (ImageJ) | (Schneider, Rasband and Eliceiri, 2012) | RRID:SCR_003070 |

### Oligonucleotide Table

| Gene | Forward primer Sequence (5'-3') | Reverse primer Sequence (5'-3') | Experiments |
|---|---|---|---|
| **qRT-PCR primers** | | | |
| **Universal Stem Loop primer (USLP)** | AAAGAAGGCGAGGAGCAGATCGAGGAAGAAGACGGAAGAATGTGCGTCTCGCCTTCTTTCNNNNNNNN | | Stem-loop qRT-PCR |
| **Universal reverse primer** | CGAGGAAGAAGACGGAAGAAT | | Stem-loop qRT-PCR |
| ***dre-miR_let7a*** | ATGCAGTGAGGTAGTAGGTTG | | Stem-loop qRT-PCR |
| ***dre-miR-92b-3p*** | ATTATTGCACTCGTCCCGGCCT | | Stem-loop qRT-PCR |
| ***dre-miR-206-3p*** | GCGTCTGGAATGTAAGGAAGTG | | Stem-loop qRT-PCR |
| ***miR_N2*** | CGCAGATCATTTTTGTGACTATG | | Stem-loop qRT-PCR |
| ***miR_N3*** | GTCAGCACTCGGACAG | | Stem-loop qRT-PCR |
| ***miR_N10*** | CGCAGTCCATCAGTCAC | | Stem-loop qRT-PCR |
| ***or42a1*** | CAGACACACACAAGGAAAACAGG | CCCCAAAACAGTCACAGCATAC | qRT-PCR |
| ***tshz1*** | ACTGCACCTCAGCAAAACAC | CGCAGTGCAGTACAGTGATG | qRT-PCR |
| ***ndr1*** | GACATCTGACCAGATATCCGCT | AAGAGTTGGATTCTCATGGCTGA | qRT-PCR |
| ***actb2*** | ATCAAGATCATTGCTCCCCCT | GGTTGGTCGTTCGTTTGAATCT | qRT-PCR |
| ***eef1a1l1*** | CACGGTGACAACATGCTGGAG | CAAGAAGAGTAGTACCGCTAGCAT | qRT-PCR |

| Gene | Forward primer Sequence (5'-3') | Reverse primer Sequence (5'-3') | Experiments |
|---|---|---|---|
| **Oligonucleotides** | | | |
| ***or42a1*_ Antisense strand** | CAGACACACACAAGGAAAACAGG | GAGAATTAACCCTCACTAAAGAAGCAACCAGTCTGTCATAAGCC | *In situ* hybridization |
| ***or42a1*_ sense strand** | GAGAATTAACCCTCACTAAAGAAGC ACAGACACACACAAGGAAAACAGG | ACCAGTCTGTCATAAGCC | *In situ* hybridization |
| **Universal T3 annealing primer** | AATTAACCCTCACTAAAGGGAGA | | miRNA mimic synthesis |
| **miRN1 mimic guide template** | AGTTGCATAGTCACAAAAATGATTCTCCCTTTAGTGAGGGTTAATT | | miRNA mimic synthesis |
| **miRN1 mimic passenger template** | ATCATTTTTGTGACTATGCAACTTCTCCCTTTAGTGAGGGTTAATT | | miRNA mimic synthesis |
| ***or42a1*-CDS mimic guide template** | TCCACATCACTCCTATGATCAGAAACATTGTCTCCCTTTAGTGAGGGTTAATT | | *or42a1*-CDS mimic synthesis |
| ***or42a1*-CDS mimic passenger template** | CAATGTTTCTGATCATAGGAGTGATGTGGATCTCCCTTTAGTGAGGGTTAATT | | *or42a1*-CDS mimic synthesis |
| **LNA Probe** | [Cy5]- AGTTGCA[+T] A[+G] [+T] [+C]AC[+A] [+A] A[+A] [+A] TGAT | | Fluorescent in situ hybridization |
